# The origin of diffuse noxious inhibitory controls

**DOI:** 10.1101/2022.03.29.486214

**Authors:** Mateusz Wojciech Kucharczyk, Francesca Di Domenico, Kirsty Bannister

**Author notes:** **Correspondence** Address: Institute of Psychiatry, Psychology and Neuroscience, Wolfson CARD, Guy’s Campus, King’s College London, London, SE1 1UL. UK.

## Abstract

Pain alerts us to actual or potential tissue damage. During acute pain, our central nervous system acts endogenously to modulate pain processing, thus reducing or enhancing pain perception. However, during chronic pain, the balance between inhibitory and facilitatory processes are tipped in favour of pro-pain modulation.

Diffuse noxious inhibitory controls (DNIC) is a naturally occurring pain inhibitory pathway that projects from the brainstem to the spinal cord to inhibit neuronal activity therein in a manner that is 1) subserved by noradrenaline, and 2) dysfunctional in chronicity. To harness its high therapeutic potential, we aimed to anatomically and functionally define DNIC.

Through employing an intersectional opto- and chemogenetic approach to modulate activity in brainstem noradrenergic nuclei, here we show that spinal neuronal firing observed upon DNIC activation during electrophysiological experiments, and animal pain thresholds observed during behavioural experiments, are modulated in a pro-pain manner upon opto-manipulation of A5 spinally projecting noradrenergic neurons, thus evidencing the DNIC origin.

Given the plasticity of the functional expression of DNIC in disease, and the success of back and forward translation of paradigms that evoke DNIC in pre-clinical and clinical models, our findings offer an attractive avenue of studies for disease specific analgesic interventions.

## Introduction

Acute pain focuses attention on a source of bodily harm. The body, possessing an endogenous ability to modulate the level of pain perceived, allows this attention to be ‘fine-tuned’ according to where it is most needed. Diffuse noxious inhibitory controls (DNIC), whereby application of a noxious ‘conditioning’ stimulus to one part of the body inhibits pain perceived in a second, remote body region, represents a form of naturally occurring pain-focusing mechanism. The analgesic DNIC phenomenon was originally described in rodents, whereby the activity of wide dynamic range (WDR) neurons is inhibited by noxious stimuli applied to various parts of the body^1^, and is observed in humans also^2^.

DNIC are pan-modal (i.e. heat and mechanical), and both conditioning and test stimuli can be of the same or different modality^3,4^. DNIC are abolished in rats following spinalisation^5^, while in humans cervical spinal cord transection^2^ or medullary retro-olive lesions (Wallenberg’s syndrome) diminishes its expression^6^. Together the data suggest a supraspinal brainstem origin for DNIC. A series of lesioning experiments previously targeted brain regions including the pontomesencephalic locus coeruleus/subcoeruleus^7^, periaqueductal grey, cuneiform nucleus and parabrachial nucleus^8^, culminating in evidence for the medullary reticular dorsal nucleus (MdD), known also as subnucleus reticularis dorsalis (SRD), as the origin of DNIC^9^. Despite these seminal findings a recent genetic, anatomically and functionally precise investigation revealed that activation of the MdD Tac1^+^ neurons facilitates thermal pain reflexes^10^. Further, in healthy rats, DNIC are shown to be sub-served by noradrenergic mechanisms mediated by activation of spinally located α_2_-adrenoceptors (α_2_-AR)^4,11,12^, and the MdD is non-catecholaminergic.

The aim of the study presented herein was to anatomically and functionally define the origin of DNIC. Noradrenergic brainstem A1-7 nuclei represent a heterogenous population of cells, whereby the discrete circuitries therein differ in topography as well as the terminal distribution of their axons^13,14^. The influence of these cells over spinal nociceptive processing is not global, and only the A5, A6 and A7 nuclei contain spinally-projecting noradrenergic neurons^15,16^. Thus, we individually modulated (activated or inhibited) A5, A6 and A7 nuclei using spatially and genetically restricted opto- and chemogenic approaches. The impact of individual nucleus manipulation on the electrophysiological properties of spinal WDR neurons and behavioural responses to mechanical and thermal stimulation, was investigated.

## Results

### Spinal α_2_-adrenoceptors mediate DNIC

The functional expression of DNIC was previously recorded in anaesthetised^4,9,17^ and wakeful animals^18^, or quantified according to markers of spinal activity^19,20^, reviewed in^21^. We examined DNIC expression in healthy rats under light isoflurane/N_2_O/O_2_ anaesthesia (slight toe pinch reflex maintained, and heart rate between 350-400 bpm). Using terminal electrophysiological recording of 95 polymodal and intensity coding (RM-ANOVA with Geisser-Greenhouse correction: [von Frey] F_(1.39, 133.40)_=377.2, P<0.0001, Tukey post-hoc) (Fig. 1a, b) lumbar deep dorsal horn WDR neurons (mean depth 854.4±6.8 μm) (Fig. S1a) we quantified functional DNIC expression as a decrease in WDR neuronal firing to a peripherally applied test stimulus upon application of a distally placed conditioning stimulus (CS). WDR von Frey-evoked firing rates were significantly decreased upon simultaneous application of CS (calibrated ear pinch) (RM-ANOVA: [DNIC] F_(1, 94)_=546.6, P<0.0001, Tukey post-hoc (Fig. 1c). Specifically, application of CS resulted in 41.5%, 49.7% and 33% inhibition of the evoked action potentials to 8 g, 26 g and 60 g von Frey application, respectively (Fig. 1c), thus representing a potent endogenous inhibitory control mechanism with over 75% of recorded units (72 of 95) achieving reduction greater than 20% for all tested forces (Fig. 1d). Spinal application of selective α_2_-AR antagonist atipamezole^22^ abolished DNIC expression (100 μg atipamezole: Two-Way RM-ANOVA: F_(1.38, 8.26)_=15.19, P<0.01, Tukey post-hoc test) (Fig. 1e, f), whereas spinal application of selective α_1_-AR antagonist prazosin^22^ failed to abolish DNIC expression (20 μg prazosin: Two-Way RM-ANOVA: F_(1.03, 5.12)_=0.57, P>0.05) (Fig. 1e, g). Neither atipamezole (Two-Way RM-ANOVA: F_(1.13, 6.78)_=0.314, P>0.05), nor prazosin (Two-Way RM-ANOVA: F_(1.19, 5.96)_=0.34, P>0.05) had any effect on basal von Frey-evoked responses (Fig. S1b, c).

**Figure 1.**
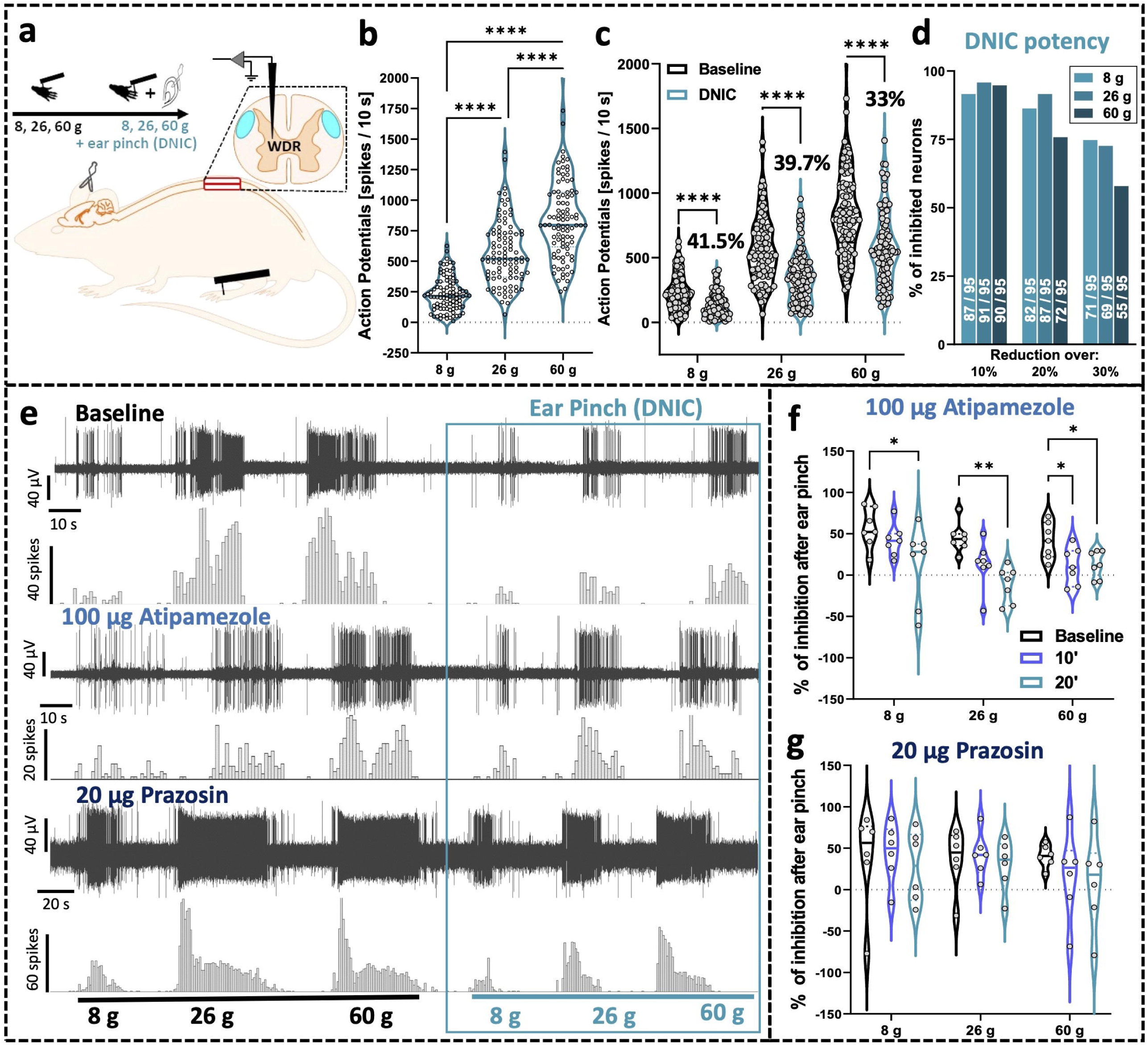
Spinal α_2_-adrenoceptors mediate DNIC. **a)** Experimental setup. **b)** Deep dorsal horn wide dynamic range (DDH-WDR) neurons code stimulus intensity (von Frey-evoked). **c)** Diffuse noxious inhibitory controls (DNIC), triggered by application of noxious ear pinch (conditioning stimulus, CS), inhibit DDH-WDR von Frey-evoked firing. **d)** Percentage of neurons inhibited by CS representing DNIC potency. Numbers overlaying bars show units which activity was reduced by a given threshold (out of 95 recorded). **e)** Example single unit DDH-WDR neuronal traces. **f)** Quantification of inhibition achieved after CS application in baseline and after α_2_-adrenoceptors block with spinal atipamezole, **g)** Quantification of inhibition achieved after CS application in baseline and after α_1_-adrenoceptors block with spinal prazosin. Data represents mean±SEM. Dots represent individual neuron studied (Baselines: N=69 rats, n=95 neurons). For pharmacology one cell was recorded per animal (atipamezole: N/n= 7, prazosin: N/n= 6). Two-way RM-ANOVA with Tukey *post-hoc*: *P<0.05, **P<0.01, ****P<0.0001.

### Inhibition of the dorsolateral funiculus abolishes DNIC

The source of spinal noradrenaline is exclusively supraspinal, and the noradrenergic fibres travel majorly via the dorsolateral funiculus (DLF)^23–25^. We probed the brain noradrenergic nuclei by retrogradely labelling descending projections to the lumbar spinal cord using spinally injected canine adenovirus (CAV) with an artificial promoter (PRS) restricting construct expression to catecholaminergic neurons (Fig. 2a-d, S1d-g). The source(s) of lumbar noradrenaline were verified by the pathways’ reconstruction using optically transparent (PACT clearing^26^) thick tissue sections (up to 1 mm) confirming primarily DLF route for the fibres (Fig. 2b; few fibres were also detected in the ventrolateral funiculi), and efficiently labelling pontine A5-A7 noradrenergic somas Fig. 2c, d, S1e). Despite unilateral virus injection in the cord parenchyma, the labelling was bilateral with a bias towards ipsilateral dominance (Fig. 2c, S1e). The noradrenergic phenotype of the labelled circuits was further confirmed by immunostaining for dopamine-β-hydroxylase (DBH) (Fig. S1f). Double-labelled neurons were identified mostly in the pontine A5, A6 and A7 nuclei ([CAV]: 24%, 12.7%, 16.6% - percentage of all DBH+ neurons therein, respectively) (Fig. S2g) corresponding with previous reports^23,24,27–29^. Using microoptrode *in vivo* electrophysiological recordings of transduced A6 neurons we also confirmed minimal light parameters required for optogenetic inhibition and activation of these neurons (Fig.1e,^12^).

**Figure 2.**
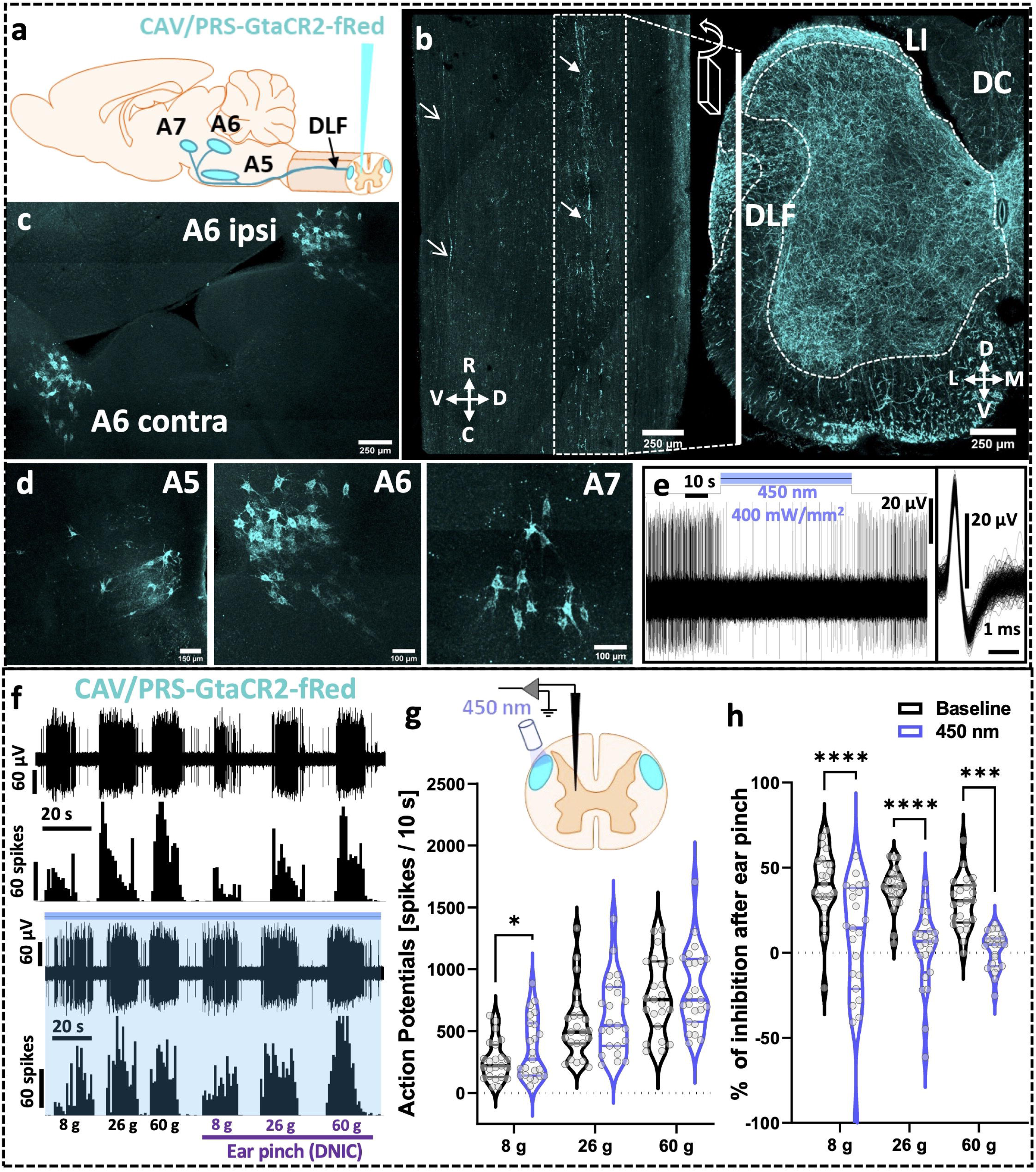
Inhibition of the dorsolateral funiculus abolishes DNIC. **a)** Experimental approach with CAV-PRS-GtACR2-fRed virus injected in the lumbar dorsal horn (DH) labels discreet brainstem noradrenergic neuronal populations (A5, A6, A7). **b)** 3D reconstruction of the light-transparent (PACT-cleared) 800 μm thick sagittal and coronal section of lumbar spinal cord evidencing labelled fibres travel via dorsolateral funiculus (DLF). Closed arrows point at DLF, open arrows point at fibres in the ventrolateral funiculus. R-rostral, C-caudal, M-medial, L-lateral, V-ventral, D-dorsal, LI-lamina I, DC-dorsal column. **c)** 3D reconstruction of the light-transparent 600 μm thick pontine coronal section showing bilateral labelling of the A6 coerulean neurons following unilateral virus injection in the lumbar DH. **d)** As in c) zoomed on the A5, A6 and A7 spinally projecting noradrenergic neurons. **e)** Representative single unit neuronal recording of the A6 GtACR2-expressing neuron inhibition following 450 nm continuous laser light illumination (400 mW/mm^2^). The inclusion shows overlay of 64 action potentials. **f)** Example traces of von Frey-evoked firing of the deep dorsal horn wide dynamic range (DDH-WDR) neurons before and after GtACR2-mediated inhibition (450 nm continuous laser light illumination, 400 mW/mm^2^, blue shaded) of the ipsilateral DLF. **g)** Noxious (26, 60 g) von Frey-evoked firing of DDH-WDR neurons is not affected by the DLF optical inhibition. **h)** Diffuse noxious inhibitory controls (DNIC), triggered by application of noxious ear pinch, are abolished after DLF GtACR2-mediated inhibition. Data represents mean±SEM. Dots represent individual neuron studied (N=18 rats, n=23 neurons). Two-way RM-ANOVA with Tukey *post-hoc*: *P<0.05, ***P<0.001, ****P<0.0001.

Next, given that most fibres travelled via the ipsilateral DLF to later bifurcate (medullary decussation), we positioned a 200 µm optic fibre directly above the DLF proximal to the recorded WDR neurons in rats expressing *Guillardia theta* anion-conducting channelrhodopsin (GtACR2)^30^, to inhibit descending noradrenergic controls with blue light (450 nm optimal for GtACR2’s activation)^31^. Given the short, transient effect of paradoxical activation upon GtACR2 axonal illumination as reported by others^30,32^, we delivered continuous 450 nm laser illumination to the DLF (400 mW/mm^2^) at least 20 s prior and throughout our sensory testing (maximum 5 minutes). Interestingly, the DLF’s optoinhibition affected only innocuous (8 g), but not noxious (26 and 60 g), von Frey-evoked basal firing of WDR neurons (Two-Way RM-ANOVA: F_(1, 22)_=4.46, P<0.05, Tukey post-hoc test: [8 g]: P<0.05, [26, 60 g]: P>0.05), suggesting the presence of a tonic noradrenergic inhibitory control restricted to innocuous mechanical stimuli (Fig. 2f, g). Importantly, DLF inhibition resulted in an almost complete reversal of the DNIC effect (Two-Way RM-ANOVA: F_(1, 22)_=80.60, P<0.0001, Tukey post-hoc test: [8, 26 g]: P<0.0001, [60 g]: P<0.001) (Fig. 2f, h).

### Inhibition of spinally projecting A5 neurons abolishes DNIC

Benefiting from the robust labelling of the A5-A7 brainstem nuclei achieved with the CAV vectors each nucleus was selectively illuminated in separate animals via a sterotaxically implanted optic fibre positioned 200 µm above the target nucleus. The use of blue light offered maintenance of selectivity in a spatially compact area (the closest distance between the nuclei is <2 mm), while sacrificing mesoscale illumination. Using this spatially and genetically restricted approach, we verified whether light-evoked inhibition of the A5, A6 or A7 nucleus, respectively, impacted basal spinal DDH-WDR neuronal firing and functional DNIC expression. Inhibition of no nuclei affected basal mechanically-evoked activity of spinal DDH-WDR neurons: (**A5**: Two-Way RM-ANOVA [450 nm] F_(1, 9)_=0.022, P>0.05; **A6**: Two-Way RM-ANOVA [450 nm] F_(1, 13)_=0.203, P>0.05; **A7**: Two-Way RM-ANOVA [450 nm] F_(1, 7)_=0.806, P>0.05) (Fig. 3a, b, d, e, g, h). Interestingly, only inhibition of spinally projecting A5 noradrenergic neurons by direct illumination of their somas abolished DNIC expression (Two-Way RM-ANOVA [DNIC] F_(1, 9)_=107.8, P<0.0001, with Tukey post-hoc: [8 g]: P<0.0001, [26, 60 g]: P<0.01) (Fig. 3a, c). Neither inhibition of the A6 nor the A7 cell groups had any effect on DNIC expression (**A6:** Two-Way RM-ANOVA [DNIC] F_(1, 13)_=0.958, P>0.05; **A7:** Two-Way RM-ANOVA [DNIC] F_(1, 7)_=0.806, P>0.05) (Fig. 3d, f, g, i).

**Figure 3.**
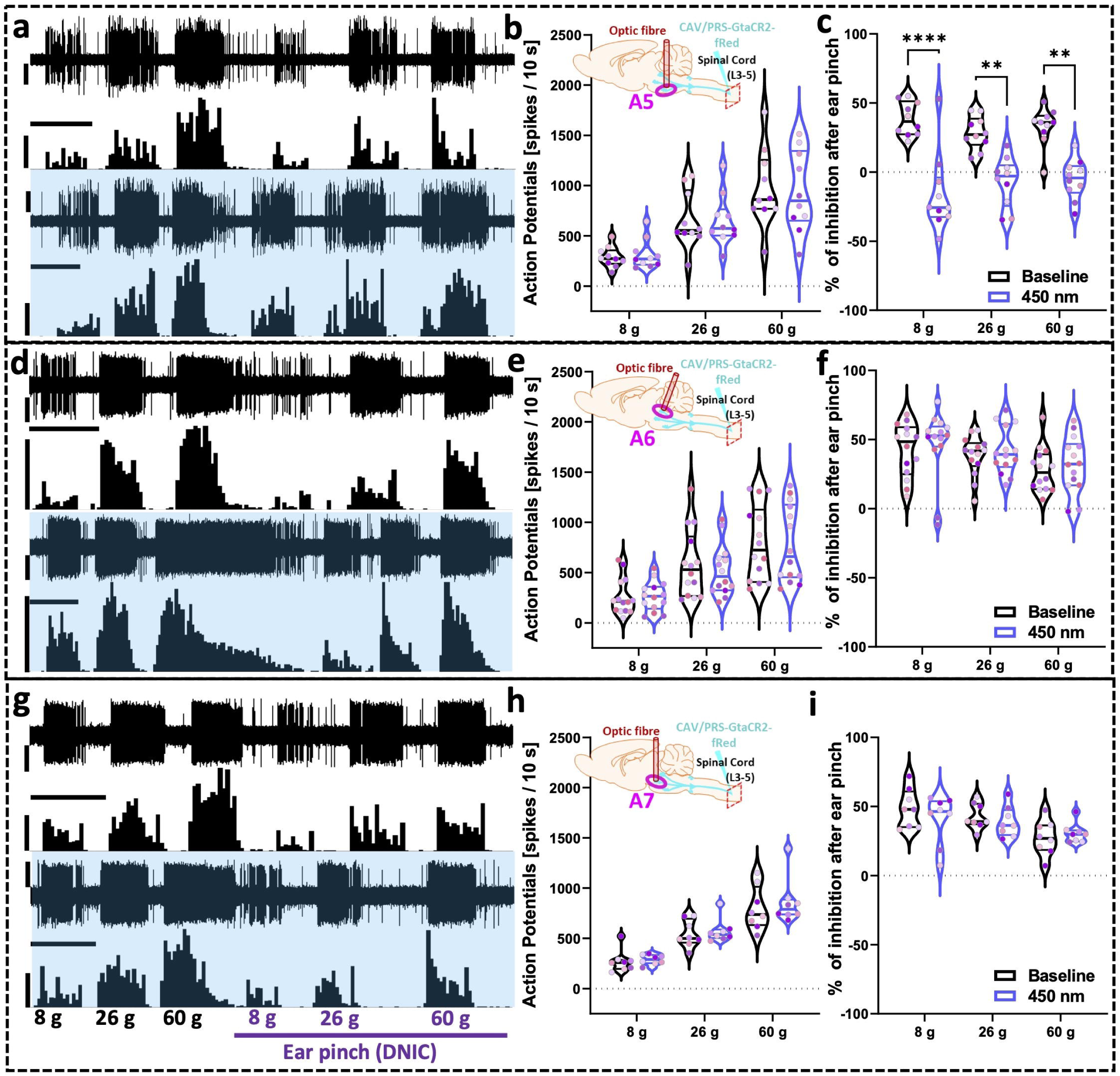
Inhibition of spinally projecting A5 neurons abolishes DNIC. **a, d, g)** Example traces of the deep dorsal horn wide dynamic range (DDH-WDR) neurons von Frey-evoked firing before and after GtACR2-mediated inhibition (450 nm continuous laser light illumination, 400 mW/mm^2^) of the labelled A5, A6, and A7 nucleus, respectively. **b, e, h)** DDH-WDR neurons basal von-Frey evoked responses are not altered optical inhibition of any of the studied A5, A6, and A7 nuclei, respectively. **c)** Diffuse noxious inhibitory controls (DNIC), triggered by application of noxious ear pinch (conditioning stimulus, CS), are abolished after A5 GtACR2-mediated inhibition, but not after A6 or A7 inhibition, **f)** and **i)**, respectively. Data represents mean±SEM. Dots represent individual neuron studied (A5: N=6 rats, n=10 neurons, A6: N=7, n=14, A7: N=6, n=8), and dots are colour coded to reflect neurons studied from the same animal. Two-way RM-ANOVA with Tukey *post-hoc*: **P<0.01, ****P<0.0001. Scale bars in a), d), g): waveform trace: 60 μV, spike count: 60 spikes, time scale: 20 s.

### The A5 neurons project directly to the spinal lamina V to mediate DNIC

Next, we implemented an intersectional labelling approach to further refine our results. We spinally microinjected AAV retrograde vectors (AAVrg,^33^) carrying floxed red-shifted cruxhalorhodopsin, Jaws^34^, fused with eGFP under pan-neuronal-specific promoter (synapsin), and a minimum of 1 week later, a second AAV9 vector encoding Cre recombinase under tyrosine hydroxylase (TH) promoter was microinjected into the relevant nuclei (A5, A6 or A7) ipsilateral to the spinal injection to restrict labelling to catecholaminergic neurons *in situ*. We found that this intersectional approach efficiently labels non-coerulean (A5 and A7) spinal noradrenergic projections but shows low efficiency for the coerulean system (A6)^35^ ([AAVrg]: 39%, 1.1%, and 12.6% - percentage of all DBH+ neurons in the A5, A6, and A7, respectively) (Fig. 4a, b, S2a, b, c). Interestingly, using this approach we did not see substantial off labelling in other, non-injected nuclei. As before, using PACT-cleared lumbar spinal cords, this intersectional approach allowed us to reconstruct the nucleus-specific projections, evidencing that the A5 fibres target directly DDH lamina IV-V (Fig. 4c), whereby WDR neurons are predominantly found (Fig. S1a), suggesting a direct A5 to DDH-WDR neuron projection.

**Figure 4.**
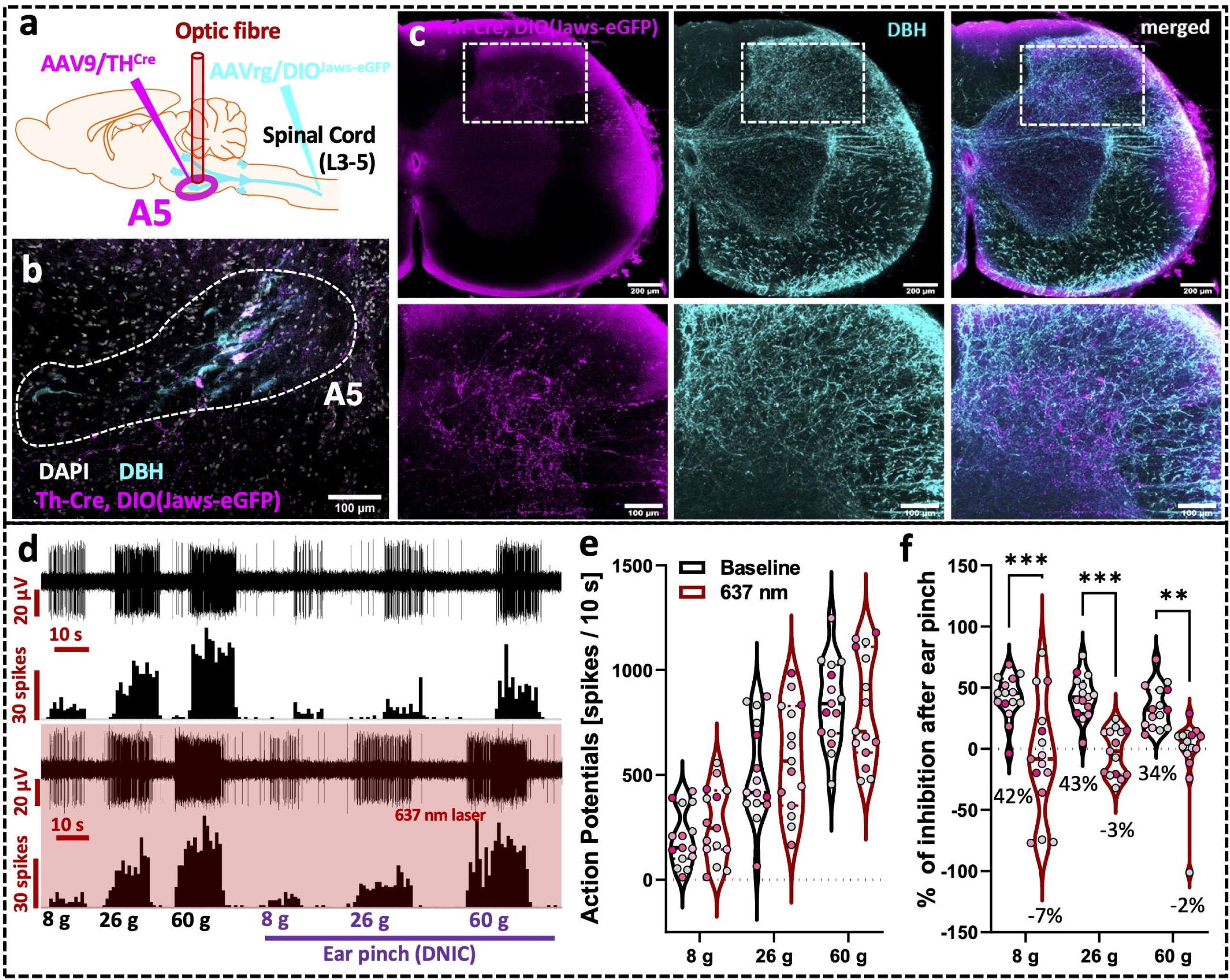
The A5 neurons project directly to the spinal lamina V to mediate DNIC. **a-b)** Experimental approach with immunohistochemical representation of the labelled nucleus. Two different adeno-associated viruses (AAV) were used to intersectionally label discreet A5 brainstem noradrenergic (dopamine-β-hydroxylase marked, DBH) neuronal population projecting to the lumbar spinal cord. **c)** Light-transparent (PACT-cleared) 800 μm thick coronal section of lumbar spinal cord evidencing accumulation of intersectionally labelled fibres in deep dorsal horn laminae IV-V. **d)** Example traces of the deep dorsal horn wide dynamic range (DDH-WDR) neurons von Frey-evoked firing before and after Jaws-mediated inhibition (637 nm continuous laser light illumination, 160 mW/mm^2^) of the labelled spinally projecting A5 neurons. **e)** DDH-WDR neurons von Frey-evoked firing is not affected by optical inhibition of the A5 nucleus. **f)** Diffuse noxious inhibitory controls (DNIC), triggered by application of noxious ear pinch (conditioning stimulus, CS), are abolished after A5 Jaws-mediated inhibition. Data represents mean±SEM. Dots represent individual neuron studied (N=10 rats, n=15 neurons), and dots are colour coded to reflect neurons studied from the same animal. Two-way RM-ANOVA with Tukey *post-hoc*: **P<0.01, ***P<0.001.

Subsequently, using this spatially and genetically restricted expression of red light (637 nm)-driven inward inhibitory chloride ion pump (Jaws), we once more verified whether light-induced inhibition of the A5, A6 or A7 nucleus, respectively, impacted spinal nociceptive processing. Despite low yeild of the A6 transduction, we performed optoinhibition with the view that higher red light (637 nm) penetration in the brain parenchyma may overcome the low number of cells labelled.

As before, no nucleus inhibition affected basal mechanically-evoked activity of spinal DDH-WDR neurons: (**A5**: Two-Way RM-ANOVA [637 nm] F_(1, 14)_=1.711, P>0.05; **A6**: Two-Way RM-ANOVA [637 nm] F_(1, 10)_=0.353, P>0.05; **A7**: Two-Way RM-ANOVA [637 nm] F_(1, 6)_=2.359, P>0.05) (Fig. 4d, e, S2d, e, g, h). Interestingly, only inhibition of spinally projecting TH+ A5 neurons by direct illumination of their somas abolished DNIC expression (Two-Way RM-ANOVA [DNIC] F_(1, 14)_=39.09, P<0.0001, with Tukey post-hoc: [8, 26 g]: P<0.001, [60 g]: P<0.01) (Fig. 4d, f). Neither inhibition of the A6, nor the A7 cell groups had any effect on DNIC expression (**A6:** Two-Way RM-ANOVA [DNIC] F_(1, 10)_=0.295, P>0.05; **A7:** Two-Way RM-ANOVA [DNIC] F_(1, 6)_=0.450, P>0.05) (Fig. S2d, f, g, i).

### Activation of spinally projecting A5 neurons mimics DNIC in the absence of a CS

Next, we adopted a previously optimised approach for CAV-mediated delivery of channelrhodopsin 2 (ChR2) to spinally projecting neurons from all three nuclei following spinal injection of CAV/PRS-ChR2-mCherry virus^12,36,37^. After confirming similar labelling pattern as for the GtACR2 constructs (Fig. 5a, S3a, b), we optoactivated spinally projecting ChR2-expressing A5 neurons, with pulsed 450 nm laser light (5 Hz, 20 ms square-wave pulses at 238 mW/mm^2^). A5 optoactivation (Two-Way RM-ANOVA [450 nm] F_(1, 14)_=7.659, P<0.05, Tukey *post-hoc* test: [8]: P>0.05, [26 g]: P<0.001, [60 g]: P<0.0001) potently inhibited mechanically evoked DDH-WDR neuron firing in the absence of CS (Fig. S3c, d), while optoactivation of the A5 nucleus had no effect on DNIC expression (**A5:** Two-Way RM-ANOVA [450 nm] F_(1, 11)_=1.02, P>0.05). The A5-mediated DDH-WDR neuronal inhibition was reversed by spinal application of 100 μg atipamezole (Two-Way RM-ANOVA [drug] F_(1.35, 6.73)_=5.36, P<0.05, Tukey *post-hoc* test: [8, 26 g]: P<0.05, [60 g]: P>0.05), confirming an α_2_-AR-mediated mechanism of DNIC expression (Fig. 5b, c).

**Figure 5.**
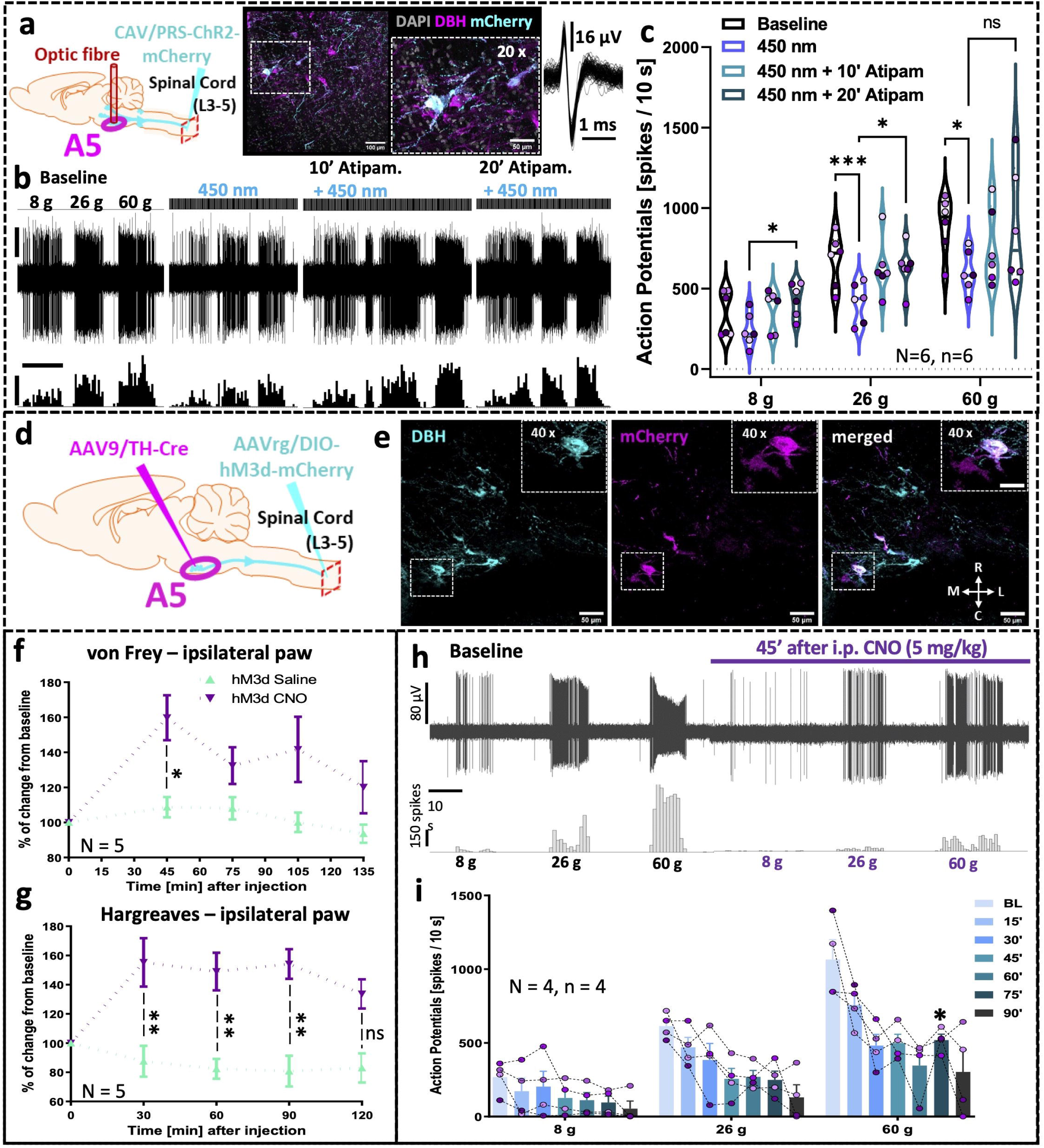
Activation of spinally projecting A5 neurons mimics DNIC in the absence of a conditioning stimulus. **a)** Experimental approach with immunohistochemical representation of the labelled A5 nucleus. CAV-PRS-ChR2-mCherry virus injected in the lumbar dorsal horn labels discreet brainstem A5 noradrenergic (dopamine-β-hydroxylase marked, DBH) neuronal populations. An inclusion shows an overlay of 50 action potentials of the neuron in b). **b, c)** Deep dorsal horn wide dynamic range (DDH-WDR) neurons von Frey-evoked firing is inhibited following optoactivation (238 mW/mm^2^ 450 nm laser light 20 ms pulses at 5 Hz) of the ipsilateral A5 nucleus that is reversible after spinal α_2_-adrenoceptors block by atipamezole. **d)** Experimental approach for intersectional labelling of the spinally projecting A5 neurons with hM3d-activatory DREADD. **e)** Immunohistochemical representation of the labelled noradrenergic (dopamine-β-hydroxylase marked, DBH) A5 nucleus. A5 chemogenetic activation (*i*.*p*. 5 mg/kg clozapine-N-oxide, CNO) induces transient elevation of **f)** mechanical and **g)** heat pain thresholds (N=5 rats). **h)** Representation of spinal single unit DDH-WDR neuron recording during hM3d-mediated activation of A5 nucleus. **i)** Quantification of neuronal responses in e). Data represents mean±SEM. Dots represent individual neuron studied (in c): N/n=6, in i): N/n=4), and dots are colour coded to reflect neurons studied from the same animal. For pharmacology one cell was recorded per animal. Two-way RM-ANOVA with Tukey *post-hoc*: *P<0.05, **P<0.01, ***P<0.001. Scale bars in b): waveform trace: 30 μV, spike count: 60 spikes, time scale: 10 s.

### Activation of the A5 nucleus mediates anti-nociception in behaving animals

The effect of activation of the A5-spinal cord pathway on antinociception was further investigated in behaving rats using a chemogenetic intersectional approach. Specifically, we employed Gq-coupled designer receptors exclusively activated by designer drugs (DREADD), hM3d, to activate the A5-spinal cord pathway in unrestrained animals (Fig. 5d, e). Corresponding with the anatomical projections of A5 nuclei we observed bilateral elevation of mechanical (Two-Way ANOVA, [time] P<0.01, F_(4, 16)_=5.260, [CNO] P<0.05, F_(1, 4)_=8.616, [Time-CNO] P>0.05, F_(4, 16)_=2.251, with Tukey *post-hoc*) (Fig. 5f) and thermal (Two-Way ANOVA, [time] P>0.05, F_(4, 16)_=2.503, [CNO] P<0.05, F_(1, 4)_=19.12, [Time-CNO] P<0.05, F_(4, 16)_=4.339, with Tukey *post-hoc*) (Fig. 5g) pain thresholds following clozapine-N-oxide injection (CNO, 5 mg/kg, *i*.*p*.). This suggest that activation of spinally projecting neurons, both in anaesthetised and in awake states, can induce antinociception, mimicking naturally evoked DNIC.

Next, 3-10 days following behavioural testing, the same animals underwent *in vivo* electrophysiology to verify whether hM3d-mediated activation of spinally projecting A5 neurons resulted in inhibition of DDH WDR neurons (4/5 rats; recording failed for one animal). Single unit extracellular recordings were made ipsilateral to the labelled A5 nucleus DDH WDR neurons (one cell per preparation) and after collection of 3 stable baseline responses activation of the A5-spinal circuity was triggered by CNO injection (5 mg/kg, i.p.). WDR neuronal activity was monitored for up to 90 minutes with 15-minute readouts (Fig. h, i). Starting from 30-minute post CNO administration, we observed a potent inhibition of DDH WDR neurons firing that lasted until the end of the recording session resembling the observed increased pain thresholds in behaving animals, with a precisely matching activity window (mean±SEM of N=4 animals per group, n=1 cell per animal; Two-Way ANOVA performed on N, [time] P<0.05, F_(2.23, 6.70)_=9.147, [von Frey] P<0.01, F_(1.22, 3.65)_=43.87, [time-von Frey] P>0.05, F_(1.86, 5.56)_=3.816, with Tukey *post-hoc*) (Fig. 5i).

## Discussion

We began our study by demonstrating, as per previous publications, that the functional expression of DNIC in healthy anaesthetised rats can be quantified, upon *in vivo* electrophysiological investigation, as a reduction in spinal WDR neuronal firing in response to a peripherally applied noxious test stimulus when a conditioning stimulus (CS) is applied concurrently in a remote body region^4,38^. The goal of the study herein was to identify the origin nucleus of the final brainstem to spinal cord projection site that leads to WDR neuronal inhibition upon application of a DNIC paradigm while also investigating whether the same nucleus to spinal cord pathway mediates anti-nociception in wakeful animals.

By performing spatially and genetically restricted optical manipulation of descending brain projections from noradrenergic A5, A6 and A7 brainstem nuclei, we demonstrated that activation of an excitatory opsin in the pontine A5 nucleus reduced the firing rate of WDR neurons in response to a peripherally applied test stimulus in the absence of CS in a manner that was reversed upon spinal application of α_2_-AR antagonist atipamezole. Conversely, upon activation of an inhibitory opsin on the same neurons, application of a CS no longer inhibited WDR firing rates in response to a peripherally applied test stimulus. Finally, chemogenetic activation of the pontine A5:SC pathway increased heat and mechanical paw withdrawal thresholds in wakeful rats. Taken together, we propose that the origin of a spinal cord projection site that governs naturally occurring analgesia is the pontine A5 noradrenergic cell group.

Despite reports of key roles for pontine cell groups A5, A6 and A7 in sympathetic activity, somatosensory transmission, and motor control respectively, that fact that all harbour the origin site of a spinally projecting noradrenergic pathway means that they must all intrinsically be capable of pain modulation via activation of spinal α-adrenoceptors (where receptor subtype and location, and pathophysiological condition, would impact the final transmission outcome)^39^. This is important to regard when considering the conserved actions of singular A5-A7 nucleus to spinal cord pathways. While some clusters of primary afferents are modality specific, superimposing the existence of different synapses on central (spinal) sites^40–42^, the ensuing action(s) of noradrenaline as released from descending pathway presynaptic terminals are likely to be rather diffuse, allowing for a broad control of multiple modalities^43–45^.

We do not predict that the DNIC pathway operates in isolation. While, anatomically, tracts within the dorsolateral funiculus are crucial for DNIC expression^25,46^, they are also crucial for ascending noxious transmission^47^; indeed it has previously been reported that, to evoke DNIC, NK1+ superficial dorsal horn projecting neurons are necessary^47–49^. Further, even though we hypothesise that the DNIC pathway is an evolutionarily conserved component of the descending pain modulatory system (DPMS), we recognise that reciprocity between the circuits that govern its expression and other modulatory controls is highly likely and, in some cases, already evidenced^12^. To fully elucidate the potential possibility, and benefit, of harnessing this DNIC as a discrete pain-inhibiting control, the nature of the 1) reciprocity and 2) precise overlap in functionality that exists between discrete A5 nucleus to spinal cord controls must be investigated. For example, mirroring that which occurs upon DNIC expression, activation of the A6 (also referred to as the locus coeruleus) was previously shown to inhibit spinal nociceptive processing^50^. As such the A6 formed a traditionally viewed inhibitory component of the DPMS, where pharmacological induction of analgesia was proposed to reflect noradrenergic actions at dorsal horn α_2_-ARs^51^, akin to the DNIC pathway’s *modus operandi*. But we now know better than to consider the sub-serving pharmacological basis for spinal neuronal inhibition upon activation of A6-SC/DNIC pathways in such simplistic terms since it undermines recent evidence of a potentiated inhibitory effect on pain-related behaviours following antagonism of spinal α_2_-ARs^36^, and opposing α_2_-AR-mediated facilitatory signalling in the brainstem^52^. The situation is complex and extends to the reciprocity of effects between neighbouring medullary nuclei. While attention and vigilance are typically associated with A6 functionality, when it comes to the immediate pain-focussing mechanism of DNIC, the A5 is crucial. This of course does not exclude a role for the A6, where activity therein seemingly alters higher brain centres in concert. Insight regarding brainstem and spinal α_2_-AR-mediated mechanisms, specifically linking DNIC attenuation to impairment of descending noradrenergic modulation from the A6 in a rodent model of joint inflammatory pain^53^, highlights the need to investigate governance of effects subsequent to A-nucleus activation in health and disease. For example, it has been previously reported that DNIC expression is abolished in rodent models of chronic pain^4,38,54,18^, highlighting the vital role that endogenous pain modulatory pathways including DNIC play in nociceptive processing/pain.

DNIC were anticipated to originate from the medullary nucleus dorsalis (MdD)^9^ and tracing studies from the MdD showed that direct projections ensue via the dorsolateral funiculus to the deep dorsal horn of the spinal cord, wherein WDR neurons reside, entirely omitting the superficial dorsal horn^55^. It is likely that MdD neurons, with their whole-body receptive field, are a vital component of an ascending relay for DNIC activation, where the MdD activates other (currently undisclosed) noradrenergic nuclei to mediate DNIC. Along with so many other aspects of DNIC governance, this requires further investigation. It is possible that the seminal lesioning studies performed^9^ corrupted facilitatory projections from the MdD to relevant brainstem noradrenergic nuclei, thus leading researchers to conclude that the MdD was the origin of DNIC. Indeed, the MdD was recently postulated to mediate facilitatory actions at the level of the spinal cord, thus lesions to this medullary region could result in baseline inhibition of spinal neurons.

Our demonstration of the impact of chemogenic activation of A5 neurons on presumed-nociceptive withdrawal thresholds in wakeful animals bears relevance for the measurement of the proposed counterpart of DNIC, the descending control of nociception (DCN^56^), in wakeful animals. It is likely that DNIC and DCN are governed differentially, wherein a higher cortical centre top-down modulatory circuitry encompasses attention to the most damaging insult in wakeful animals^56^. Nonetheless, we have indicated that the origin of the final spinal cord output of a noradrenergic pathway that governs inhibition of spinal nociceptive processing upon CS application in anaesthetised versus wakeful animals, is the same. Consideration of the processes that underlie endogenous pain inhibition upon conditioning in behaving rodents has included age and sex as well as hormonal and net descending facilitatory control influences^18,57,58^, where DCN likely involves DNIC mechanisms^56^. Incorporating higher cognitive and affective influences, DNIC are termed ‘conditioned pain modulation’ (CPM) in humans^59,60^ and it is noteworthy that, even when recorded in anaesthetised rodents, the functional expression of DNIC is influenced by subcortical brain regions associated with emotional processing^18^. Pharmacotherapies that modulate noradrenergic transmission reinstate dysfunctional CPM in certain chronic pain patients^61–63^, and this back translates to rodent studies^4,38,64^. Interestingly, the MdD is modulated by higher brain centres including the neocortex^65^, linking the analgesic actions of distraction, and pointing at the relationship with cognitive processes.

## Conclusions

The innate ability of the body to modulate the level of pain perceived following, for example, a noxious insult at the periphery reflects the activation of modulatory circuits that can reduce or facilitate the spinal neuronal response. Thus, the final pain percept depends on the context of the insult. If two noxious stimuli are delivered concurrently to distant body regions, activation of DNIC serves to focus attention, preserving bodily integrity. The temporal aspect of DNIC means that the pain inhibits pain phenomenon is somehow inhibited, or dysfunctional, when the body experiences persistent pain. This requires further investigation, i.e. how long is too long in terms of the presence of pain for the DNIC circuit to stop working? The input threshold of the test stimulus and CS that is required (minimal and maximal) to induce DNIC expression is also unknown.

The experiments described in this study highlight a key role for A5 spinal-cord projecting neurons in endogenous pain inhibition, thus we provide a solid foundation for future key studies that aim to harness the body’s endogenous ability to reduce pain.

## Author contributions

M.W.K and K.B. conceptualised the study. M.W.K. designed the methodology. M.W.K. performed physiological and PACT experiments. F.D.D. performed immunohistochemical analysis. M.W.K. performed formal analysis and data visualisation. M.W.K. and K.B. wrote the manuscript. K.B. acquired the funding and administered the project. All authors have read and agreed to the published version of the manuscript.

## Acknowledgements

Authors would like to thank Professor Anthony Pickering for supplying us with CAV virus used in this study and Professor Stephen McMahon for start-up equipment funding.

## Supplementary figure legends

**Figure S1. Spinal α**_**2**_**-adrenoceptors mediate DNIC. a)** Lumbar spinal cord electrode depths of all the recorded deep dorsal horn wide dynamic range (DDH-WDR) neurons. **b)** Quantification of von Frey-evoked action potentials before and after α_2_-adrenoceptors block with spinal atipamezole, **c)** Quantification of von Frey-evoked action potentials before and after α_1_-adrenoceptors block with spinal prazosin. **d)** Experimental approach with CAV-PRS-GtACR2-fRed virus injected in the lumbar dorsal horn (DH) labels discreet brainstem noradrenergic neuronal populations (A5, A6, A7). **e)** 3D reconstruction of the light-transparent (PACT-cleared) 600 μm thick pontine coronal section showing bilateral labelling of the A6 and A5 neurons following unilateral virus injection in the lumbar dorsal horn. **f)** Immunohistochemical representation of the noradrenergic (dopamine-β-hydroxylase marked, DBH) labelled A5, A6 and A7 nuclei after CAV-PRS-GtACR2-fRed virus injected in the lumbar dorsal horn. **g)** Quantification of the CAV labelling efficiency shown as a percentage of all DBH+ neurons in a given ipsilateral to the virus injection nucleus (i-ipsi, c-contra). Data represents mean±SEM. For electrophysiology, dots represent individual neuron studied (all recorded neuron depths: N=69 rats, n=95 neurons. For pharmacology one cell was recorded per animal (atipamezole: N/n= 7, prazosin: N/n= 6), and dots are colour coded to reflect neurons studied from the same animal. Two-way RM-ANOVA. For histochemical quantification dots represent individual animal data as a mean from 6-8 brain slices per rat.

**Figure S2. Intersectional viral strategy for selective labelling of discreet spinally projecting noradrenergic brainstem nuclei. a, b)** Experimental approach with immunohistochemical representation of the labelled A6 and A7 nucleus. Two different adeno-associated viruses (AAV) were used to intersectionally label discreet brainstem noradrenergic (dopamine-β-hydroxylase marked, DBH) neuronal populations projecting to the lumbar spinal cord (A6, A7). **c)** Percentage of double labelled DBH neurons expressing viral tag (eGFP) in A5-A7 nuclei after ipsilateral injection of AAV viruses. Note the low labelling efficiency for A6 coerulean neurons (<2%). **d, g)** Example traces of the deep dorsal horn wide dynamic range (DDH-WDR) neurons von Frey-evoked firing before and after Jaws-mediated inhibition (637 nm continuous laser light illumination, 160 mW/mm^2^) of the labelled spinally projecting A6 and A7 neurons, respectively. **e, h)** DDH-WDR neurons von Frey-evoked firing is not affected by optical inhibition of the A6 or A7 nucleus, respectively. **f, i)** Diffuse noxious inhibitory controls (DNIC), triggered by application of noxious ear pinch (conditioning stimulus, CS), are not affected after A6 or A7 Jaws-mediated inhibition. respecitvely. Data represents mean±SEM. Dots represent individual neuron studied (A6: N=7 rats, n=11 neurons, A7: N=6 rats, n=7 neurons), and dots are colour coded to reflect neurons studied from the same animal. For histochemical quantification dots represent individual animal data as a mean from 6-8 brain slices per rat. Two-way RM-ANOVA.

**Figure S3. Optogenetic activation of the spinally projecting A5 neurons inhibits spinal neuron activity. a, b)** Experimental approach with immunohistochemical representation of the labelled A5 nucleus. CAV-PRS-ChR2-mCherry virus injected in the lumbar dorsal horn labels discreet A5 brainstem noradrenergic (dopamine-β-hydroxylase marked, DBH) neuronal population **c)** Deep dorsal horn wide dynamic range (DDH-WDR) neurons von Frey-evoked firing is inhibited following optoactivation (238 mW/mm^2^ 450 nm laser light 20 ms pulses at 5 Hz) of ipsilateral spinally projecting A5 neurons. **d)** A5 optoactivation does not abolish diffuse noxious inhibitory controls (DNIC), triggered by application of noxious ear pinch, expression. Data represents mean±SEM. Dots represent individual neuron studied (N=7 rats, n=12 cells), and dots are colour coded to reflect neurons studied from the same animal. Two-way RM-ANOVA with Tukey *post-hoc*: ***P<0.001, ****P<0.0001.

## Materials and Methods

### Animals

Male Sprague-Dawley rats (Envigo, UK) were used for experiments. Animals were group housed on a 12:12 h light–dark cycle. Food and water were available *ad libitum*. Animal house conditions were strictly controlled, maintaining stable levels of humidity (40–50%) and temperature (22 ± 2°C). All procedures described were approved by the Home Office and adhered to the Animals (Scientific Procedures) Act 1986. Every effort was made to reduce animal suffering and the number of animals used was in accordance with International Association for Study of Pain (IASP)^66^ and ARRIVE ethical guidelines^67^.

In this study we used 70 rats, assigned to the groups as follows: SC-CAV/PRS-GtACR2 injected: 6 rats for A5, 7 for A6, 6 for A7; SC-CAV/PRS-ChR2 injected: 10 rats for A5; intersectional AAV approach: (Jaws) A5-10 rats, A6-7 rats, A7-6 rats, and hM3d (5 rats); plus 13 naïve rats for basal pharmacology. In total 95 DDH WDR neurons were recorded from 69 rats in 117 experimental approaches as listed in the supplementary data spreadsheet.

### Virus injections

#### Spinal cord injections

Rats weighing 60-70 g were anaesthetised using isoflurane (3–5% for induction and 1.5–2% for maintenance in 1 l/min oxygen flow, Piramal, UK) and maintained at around 37°C using a homeothermic heating mat and 50 μl of Meloxicam (2 mg/kg, Metacam®, Boehringer Ingelheim, Berkshire, UK) was subcutaneously administered for post-operative pain management. Animals were fixed in a stereotaxic apparatus (Kopf Instruments, UK), their lumbar region was clamped and T12-L1 intervertebral space was exposed by bending the lumbar the region rostrally providing easy access to the underlaying dura and L3-4 spinal cord without the need for extensive laminectomy.

For intersectional approach (paired with the brainstem injection, see next section): bilaterally, two paired injections (200 μm lateral from the midline and 750 μm apart rostro-caudally; first pair at 850 and second pair at 450 μm below the L3-L4 cord surface) of AAVrg viral particles with improved retrograde axonal transport^33^ were made to transduce descending spinal projections. Following AAVs were used: AAVrg-CAG-FLEX-rc[Jaws-KGC-GFP-ER2] (titer >7×10^12^ vg/ml, viral prep #84445-AAVrg, Addgene, US,^34^) or AAVrg-hSyn-DIO-hM3D(Gq)-mCherry (titer >7×10^12^ vg/ml, viral prep #44361-AAVrg, Addgene, US,^68^). For canine adenovirus (CAV) global transduction of descending projections, three unilateral lumbar dorsal horn injections were performed (200 μm lateral from the midline and 750 μm apart rostro-caudally; first at 850, second at 650, and third at 450 μm below the L3-L4 cord surface). For optogenetic activation CAV encoding for channelrhodopsin 2 under the control of catecholamine-specific synthetic promoter (sPRS, no brain injection required, see below) was used (CAV/sPRS-hChR2(H134R)-mCherry, titer >3×10^10^ TU/ml, PVM, Montpellier, a gift from Professor Anthony Pickering, University of Bristol^36,37^), while for inhibition CAV encoding for *Guillardia theta* anion-conducting channelrhodopsin (stGtACR2)^30,31^ under PRS promoter was used (CAV/PRS-stGtACR2-FRed, titer >8×10^10^ TU/ml, PVM, Montpellier). Injections were made with a glass pulled micropipette (10-20 μm tip) coupled to electronically controlled nanoinjector (Nanoliter 2010, WPI, FL, US) facilitating precise delivery with minimal damage. Each injection was of 400 nl with 2 nl/s delivery rate and minimal 3-5 minutes wait between slow pipette retraction. The micropipettes were filled with inert mineral oil (extrusion medium), and the virus-oil interface was monitored to ensure injection. The wound was irrigated with saline and the incision was closed with wound clips and postsurgical glue (Vetabond, 3M, UK).

### Brainstem injections

Brainstem injections were made on rats weighting 190-210 g (around 2 weeks after spinal cord surgery). Animals were anesthetized with i.p. ketamine (5 mg/100 g, Vetalar; Pharmacia) and medetomidine (30 µg/100 g, Dormitor; Pfizer) until loss of paw withdrawal reflex and perioperative analgesia was achieved by the s.c. injections of meloxicam (2 mg/kg, Metacam^®^, Boehringer Ingelheim, Berkshire, UK). The rat was placed in a high precision stereotaxic frame (Kopf Instruments, UK) and core temperature was maintained at 37°C using a homeothermic blanket (Harvard Apparatus, US). Aseptic surgical techniques were used throughout. Using a 0.7 mm dental drill a hole was made in the skull directly above the targeted structure. The following coordinates were used: **A6 (locus coeruleus)**: 10° rostral angulation (to avoid puncturing the sinus), from lambda: RC: -2.1 mm, ML: +1.3 mm, and three injections at -6.2, -6.0, -5.8 mm deep from the cerebellar surface (2 mm deeper from lambda); **A5:** no angulation, from lambda: RC: -0.8 mm, ML: +2.4 mm, and three injections at -9.8, -9.6, -9.4 mm deep from lambda, **A7:** no angulation, from lambda: RC: +0.1 mm, ML: +2.8 mm, and three injections at -7.8, -7.6, -7.4 mm deep from lambda. Three injections of AAV9-rTH-PI-Cre-SV40 (functional, titer >7×10^12^ vg/ml, viral prep #107788-AAV9, Addgene, US, a gift from James M. Wilson) or were made analogously to spinal injections. Each injection was of 300 nl, every 200 μm starting from deepest point chosen (DV) with 2 nl/s delivery rate and minimal 2-3 minutes between slow pipette retraction. 5 minutes after final injection the micropipette was retracted over the course of 4-5 minutes. The micropipettes were filled with inert mineral oil (extrusion medium) and the virus-oil interface was monitored to ensure injection. The wound was irrigated with saline and closed with Vicryl 4-0 absorbable sutures and wound glue (VetBond 3M, UK). Anaesthesia was reversed with s.c. injection of atipamezole (Antisedan, 0.1 mg/100 g, i.p.; Pfizer). The animals were placed in a thermoregulated recovery box until fully awake. Two to three weeks were allowed for the transgene(s) expression.

### Spinal Cord In Vivo Electrophysiology

*In vivo* electrophysiology was performed on animals weighing 240–300 g as previously described^69^. Briefly, after the induction of anaesthesia, a tracheotomy was performed, and the rat was maintained with 1.5% of isoflurane in a gaseous mix of N_2_O (66%) and O_2_ (33%). Core body temperature was monitored and maintained at 37 °C by a heating blanket unit with differential rectal probe system. Electrocardiogram (ECG) was monitored by two intradermal needles inserted in front limbs with signal amplified by the Neurolog system consisting of AC preamplifier (Neurolog NL104, gain x200), through filters (NL125, bandwidth 300 Hz to 5 KHz) and a second-stage amplifier (Neurolog NL106, variable gain 600 to 800) to an analogue-to-digital converter (Power 1401 625kHz, CED). Craniotomy was performed to gain stereotaxic access to the ipsilateral LC for either optic fibre or micropipette insertion as described in following sections. A laminectomy was performed to expose the L3–L5 segments of the spinal cord, the cord was clamped to minimise movement, dura was carefully removed with the aid of surgical microscope, and the recording area was secured by saline-filled well made in solidified 2% low melting point agarose (made in saline also). Using a parylene-coated, tungsten electrode (125 μm diameter, 2 MΩ impedance, A-M Systems, Sequim, WA, USA), wide dynamic range neurons in deep laminae IV/V (∼650–900 μm from the dorsal surface of the cord) receiving intensity-coding afferent A-fibre and C-fibre input from the hind paw were sought by periodic light tapping of the glabrous surface of the hind paw. Extracellular recordings made from single neurones were visualized on an oscilloscope and discriminated on a spike amplitude and waveform basis. Specifically, the signal from the electrode’s tip was processed via headstage connected to the neurolog system consisting of AC preamplifier (Neurolog NL104, gain x200), through HumBag (Quest Scientific, North Vancouver, Canada) used to remove low frequency noise (50–60 Hz), via a second-stage amplifier (Neurolog NL106, variable gain 600 to 800), filters (NL125, bandwidth 1000 Hz to 5 KHz) and spike-trigger (Neurolog NL106, variable gain 600 to 800) to an analogue-to-digital converter (Power 1401 625kHz, CED). Spike trigger was visualised on a second oscilloscope channel and manually set to follow single unit spikes. Its analogue signal was digitalised via event input to build stimulus histogram in real time along the waveform recordings. All the data were captured by an analogue-to-digital converter (Power 1401 625kHz, CED) connected to a PC running Spike2 v8.02 software (Cambridge Electronic Design, Cambridge, UK)) for data acquisition, analysis and storage. Simultaneous ECG monitoring and transistor–transistor logic (TTL) triggers (i.e. for the lasers, see below) were additionally coupled to Spike 2 recording traces via CED-1401 analogue inputs.

### Stimulation paradigm in all electrophysiological recordings

Natural mechanical stimuli, including von Frey filaments (8 g, 26 g and 60 g) and von Frey filaments with concurrent ipsilateral noxious ear pinch (15.75 × 2.3 mm Bulldog Serrefine ear clip; InterFocus, Linton, United Kingdom), were applied in this order to the receptive field for 10 s per stimulus. The noxious ear pinch was used as a conditioning stimulus (CS) to trigger diffuse noxious inhibitory control (DNIC,^4,12,54^) and was quantified as an inhibitory effect on neuronal firing during ear pinch to its immediate respective von Frey filament applied without the conditioning stimulus (% of inhibition after ear pinch). A minimum 30 s non-stimulation recovery period was allowed between each test in the trial. A 10-minute non-stimulation recovery period was allowed before the entire process was repeated for control trial number 2 and 3. The procedure was repeated 3 times and averaged only when all neurons met the inclusion criteria of 10% variation in action potential firing for all mechanically evoked neuronal responses. No animals were excluded from analysis.

### In vivo spinal pharmacology with electrophysiological monitoring

After collection of predrug baseline control data as outlined above, atipamezole (a α_2_-AR antagonist: 100 μg; Sigma-Aldrich, Gillingham, United Kingdom, dissolved in 97% normal saline, 2% Cremophor [Sigma, UK], 1% dimethyl sulfoxide [DMSO; Sigma, UK] vehicle), prazosin hydrochloride (α_1_-AR antagonist: 20 μg, Sigma-Aldrich, Gillingham, United Kingdom, dissolved in water for injections) was administered topically to the spinal cord in 50 μl volumes following gentle removal of residing saline in the agarose well. Clozapine-N-oxide (CNO, 5 mg/kg in saline) was injected intraperitoneally. Each individual drug dose effect (one stable neuron assessed per rat) was followed for up to 40 minutes with tests performed typically at 3 time points (starting at 10, 20 and 30 minutes), except for CNO, which was monitored up to 90 minutes with 15 minutes intervals. For each time point, a trial consisted of consecutive stable responses to von Frey and DNIC (von Frey with concurrent ipsilateral ear pinch).

### Optogenetics

#### Light stimulation during spinal WDR recordings

The 450 nm laser (Doric Lenses, Quebec, Canada) or 637 nm laser (Thorlabs, UK) was externally TTL-triggered by the neurolog system (NeuroLog system, Digitimer, UK) to deliver defined light pulses (238 mW/mm^2^, 20 ms pulse width at 5 Hz for 450 nm laser pulses to activate ChR2) or continuous illumination (at 160 mW/mm^2^ for 637 nm laser for Jaws, or at 400 mW/mm^2^ for 450 nm laser for GtACR2). In brief, the laser light was coupled to a multimode 200 um patch cord (0.39 NA, #M75L01, Thorlabs, UK) and via SMA to SMA mating sleeve (#ADASMA, Thorlabs, UK) to second ferrule-terminating multimode 200 μm patch cord (0.39 NA, #M77L01, Thorlabs, UK) directly interconnected to multimode stainless steel 20 mm long cannula (200 μm diameter, 0.39 NA, #CFM12L20, Thorlabs, UK). The power density was adjusted for each preparation (measured using PM16-130 power meter (Thorlabs, UK) at the tip of implantable 200 μm fibre^36^. After desired power was achieved the fibre was slowly inserted in the brainstem nucleus ipsilateral to the injected vectors and the recorded spinal WDR neurons. The fibre was lowered using precise hydraulic micromanipulator (Narishige, Japan) mounted on stereotaxic frame (Kopf) adjusted to desired coordinates to target studied nucleus (same coordinates were used as in the brainstem virus injections). Spinal WDR neurons were characterised by three stable baseline responses followed by three optically modulated responses. For combined optogenetics and spinal pharmacology, after collecting three stable baseline and three stable optoactivation responses (averaged, if stable), a drug (100 µg atipamezole) was applied topically on the exposed spinal cord surface, right above the recording site. To assess simultaneous action of the drug and the A5 optoactivation, light pulses were delivered 30 s before and throughout each series of tests (approximately 5 minutes per series) and minimally 5 minutes of the recovery time was allowed between the tests. Pharmacology was monitored every 10 minutes for 30-40 minutes (each test with optoactivation). At the end of every experiment, animals were sacrificed by the overdose of isoflurane and transcardially perfused with cold saline followed by 4% paraformaldehyde for anatomical evaluation.

### A6 neuron recording and optoinhibition

A simultaneous recording and optical stimulation of the transduced A6 neurons were made using microoptrodes as described earlier with minor modifications^70^. For the A6 recordings and optoactivation, the all-glass recording microptrode with 20 µm tip diameter consisted of the recording core filled with 3 M sodium acetate (resulting in 2-3 MΩ resistance) and the parallel gradient index (GRIN) optical core (a gift from Professor Yves De Koninck, Laval University, Canada) coupled to the optic fibre (multimodal, 200 µm core diameter, 0.39 NA, #M77L01, Thorlabs, UK) was used. The GtACR2-expressing A6 neurons were optoinhibited by 450 nm laser (Doric Lenses, Quebec, Canada) light continuous illumination as described above.

### Behavioural testing with DREADD

Clozapine-N-oxide (CNO, Cayman, UK) was used to activate virally-delivered hM3d DREADD restricted to spinally projecting A5 catecholaminergic neurons (AAV9/TH.Cre). Before experiments 25 mg CNO was dissolved in 300 μl of pure dimethyl sulfoxide [DMSO; Sigma, UK] and 30 μl aliquots were kept frozen until use. On the experimental day, an aliquot was prewarmed to room temperature, vortexed and 470 μl of sterile saline was added resulting in 2.5 mg/0.5 ml ready to use CNO solution. Recognising CNO’s reversed metabolism, we have restricted our behavioural analysis to the first 2 hours from administration. A day before pharmacological experiments animals were habituated in the testing conditions for the same time as that of the experimental procedure (c.a. 4 hours). On the first experimental day all animals received i.p. vehicle (saline with DMSO) injection of the same volume as would be use for the CNO solution. Two days after collection of saline control responses, rats received i.p. CNO (5 mg/kg) injection and their thermal and mechanical thresholds were retested. Each testing was preceded by a 60 min acclimatisation period, during which rats spent 30 minutes exploring each Hargreaves and von Frey compartments.

### Hargreaves test

Testing compartments consisted of 20×20×25 cm plexiglass boxes with glass floor. Heat thresholds were assessed by application of infrared beam (Hugo Basile, Italy) to the plantar surface proximal to the digits of the ipsilateral and contralateral hind paws. The IR beam was applied 3 times and withdrawal responses elicited were automatically detected by the device and time to withdrawal was taken as a measure. Data was collected at 0, 30, 60, 90 and 120 minutes from the drug/vehicle injection. Results are presented as a mean ± SEM.

### Von Frey test

Testing compartments consisted of 20×20×25 cm plexiglass boxes with wire grid floor. Mechanical thresholds were assessed by application of automatic von Frey filaments with 26 g cut off (Hugo Basile, Italy). The testing filament was applied 3 times to the plantar surface proximal to the digits of the ipsilateral and contralateral hind paws. Withdrawal responses elicited by the filament were automatically detected by the device and reflective mass applied taken. Data was collected at 0, 45, 75, 85 and 135 minutes from the drug/vehicle injection. Results are presented as a mean ± SEM.

### Immunohistochemistry

Animals were sacrificed by the overdose of anaesthetic and transcardially perfused with cold phosphate buffer saline (PBS) followed by 4% paraformaldehyde (PFA) in phosphate buffer (pH 7.5). Next, collected spinal cords and brains were post-fixed in 4% PFA for 3-4 days at 4°C, followed by 3-4 days at 4°C in 30% sucrose. Once tissue density equilibrated, lumbar spinal cords and brains were precut into 5 mm thick coronal fragments with razor blades and the aid of rat brain matrix. Obtained fragments transferred to optimum cutting temperature (OCT)-filled moulds were snap-frozen in liquid nitrogen and stored frozen until further analysis. Next the OTC embedded tissue was cryo-sectioned (Bright Instruments, UK) to 25 μm thick coronal slices subsequently collected on eight Menzel-Gläser Superfrost Plus slides (a slice collected every 200 μm) and stored in -20°C until staining. Once dried (45°C for an hour) and briefly washed with 50% ethanol, sections were outlined with a hydrophobic marker (PAP pen, Japan), rehydrated and blocked with 10% donkey serum in blocking solution (0.03% NaN_3_, 0.3% Triton X-100 in PBS, pH=7.5) for two hours prior to overnight incubation at room temperature with primary antibodies against dopamine-β-hydroxylase (DBH, a marker of noradrenergic neurons: Mouse, 1:500, Millipore, MAB308, UK), mCherry (Rabbit, 1:500, Abcam, ab167453, UK), fRed (rabbit anti-tRFP, 1:500; AB233, Evrogen), or eGFP (chicken, 1:1000, ab13970; Abcam, United Kingdom). Slides were then PBS washed and incubated with the appropriate fluorophore-conjugated secondary antibodies in blocking solution (Donkey anti-Rabbit, Alexa Fluor 568, A10042, Invitrogen, Eugene, OR, US; Donkey anti-Mouse, AlexaFluor 488, A21202, Invitrogen, Eugene, OR, US; all used at 1:1000 dilution) for 4 hours to overnight at room temperature. Slides were protected with mounting media (Fluoromount-G with DAPI, eBioscience, UK) and coverslips and stored in darkness at 4°C until imaging.

Samples were typically imaged with an LSM 710 laser-scanning confocal microscope (Zeiss) using Zeiss Plan Achromat 10x (0.3 NA) and 20 x (0.8 NA) dry objectives and analysed with Fiji Win 64. For quantification, samples were imaged with 20x dry objective on Zeiss Imager Z1 microscope coupled with AxioCam MRm CCD camera. The acquisition of images was made in multidimensional mode and the MosaiX function was used to construct the full view. 6-8 slices were imaged per animal. Cell counting was carried out on the Fiji Win 64 utilising cell counter plugin. On average, 20-30 brainstem sections were imaged for quantification.

### Passive Tissue Clearing (PACT)

A passive CLARITY tissue clearing technique (PACT) (detailed in^26^) has been implemented to allow thick >1000 µm tissue fragments imaging in CAV/PRS-GtACR2-fRed injected rats. Briefly, following transcardial perfusion of deeply anaesthetised rats with a cold PBS and a cold 4% PFA in phosphate buffer, pH=7.5, spinal cords and brains were extracted and post-fixed in 4% PFA for 3-4 days in 4°C. After fixation, samples were pre-cut using vibratome at 600 μm coronal brainstem sections, or 800 μm coronal or sagittal spinal cord sections. Slices were then transferred directly to ice-cold A4P0 solution consisting of: 4% acrylamide monomer (40% acrylamide solution, cat. 161-0140, Bio-Rad, UK), 0.25% VA-044 (thermoinitiator, Wako, US) in 0.01 M PBS, pH=7.4, and incubated at 4°C overnight in distilled water prewashed (to remove anticoagulant) vacutainer tubes (Vacutainer, #454087, Greiner GmbH, Austria). The next day, samples were degassed by piercing the septum with a 20G needle connected to a custom-build vacuum line. The residual oxygen was replaced with nitrogen by 2 min bubbling of the solution with pure nitrogen (BOC, UK) via a long, bottom-reaching 20G needle, and a second short needle pierced to allow gases to exhaust. Throughout, samples were kept on ice to prevent heating and consequent premature A4P0 polymerisation. After achieving oxygen-free conditions, samples were polymerised by 3 h incubation in a 37°C water bath. Following polymerisation, the excess honey-like polyacrylamide gel was removed with tissue paper, and samples were transferred to 50 ml falcon tubes filled with clearing solution. 10% SDS (#L3771, Sigma-Aldrich, UK) in PBS, pH=8.0, was used for passive clearing. Samples were incubated on a rotary shaker at 37°C and 75 rpm (Phoenix Instruments, UK) until reaching the appropriate transparency (usually 3-4 days). Next, all samples were washed extensively with PBS, pH=7.5, on rotary shaker at room temperature, by replacing the solution 4-5 times throughout the course of 1 day to remove the SDS with extracted lipids. Following washing, samples were treated with primary antibody in blocking buffer consisting of 2% normal donkey serum in 0.1% Triton X-100 in PBS, pH=7.5 with 0.03% sodium azide. 500-1000 μl of rabbit anti fRed (1:500; AB233, Evrogen) primary antibody was used per slice in a 2 ml Eppendorf tube. Samples were incubated with the primary antibody at room temperature, with gentle shaking for 3 days. This was followed by 4-5 washing steps with PBS over the course of a day days. Next, the samples were incubated with gentle agitation, at room temperature, in darkness, for 3 days with the goat anti-rabbit (Alexa Fluor 647, A21244, Invitrogen, Eugene, OR, US) fluorophore-conjugated secondary antibody (1:500 in the blocking buffer). Thereafter, samples were washed extensively with PBS at least 5 times over 1-2 days at room temperature. Finally, samples were overnight incubated in the refractive index-matching solution (RIMS, refractive index = 1.47) consisting of 40 g of Histodenz (#D2158, Sigma-Aldrich, UK) dissolved in 30 ml of PBS, pH=7.5 with 0.03% sodium azide. 400-600 μl of RIMS was used per structure. Samples were placed in fresh RIMS in custom-made glass slide chambers, covered with coverslips, and equilibrated for few hours before imaging.

Samples were imaged with a Zeiss LSM 780 one-photon confocal upright microscope, equipped with EC Plan-Neofluar 10x 0.3 NA, Ph1 dry objective (WD=5.3 mm, cat. 420341-9911, Zeiss, Germany) and 633 nm laser lines. Scans were taken with 2048×2048 pixel resolution, with 4-5 μm optical section typically spanning 400-700 μm of scanned depth (resulting in 100-150 planes) with auto Z-brightness correction to ensure uniform signal intensity throughout the sample. Images were exported from Zen 2012 Blue Edition software (Carl Zeiss Microscopy GmbH, Germany). Next graphical representations, 3D-rendering, animations, maximal intensity projections within selected z-stacks and further analysis were obtained with open-source Fiji (ImageJ) equipped with appropriate plugins.

### Quantification and Statistical Analysis

Statistical analyses were performed using SPSS v25 (IBM, Armonk, NY, USA). All data plotted in represent mean ± SEM. Typically, up to 4 WDR neurons were characterised per preparation (n), and data were collected from at least 5 rats per group (N). Single pharmacological investigation was performed on one neuron per animal. Statistical analysis was performed either on number of neurons (n) for populational studies or number of animals (N) for pharmacological studies. Therefore, throughout the manuscript “n” refers to the number of cells tested and “N” to the number of animals tested. Detailed description of the number of samples analysed and their meanings, together with values obtained from statistical tests, can be found in each figure legend. Symbols denoting statistically significant differences were also explained in each figure legend. Main effects from analysis of variance (ANOVA) are expressed as an F-statistic and *p*-value within brackets. Throughout, a p-value below 0.05 was considered significant. Uncorrected two-way repeated-measures (RM) ANOVA with the Tukey post-hoc was used to assess von Frey and DNIC responses in the baseline conditions. For pharmacological experiments, Geisser-Greenhouse correction was used for RM-ANOVA. GraphPad Prism was used to analyse the data.

## References

1. Le Bars, D., Dickenson, A. H. & Besson, J. M. Diffuse noxious inhibitory controls (DNIC). I. Effects on dorsal horn convergent neurones in the rat. Pain 6, 283–304 (1979).

2. Roby-Brami, A., Bussel, B., Willer, J. C. & Lebars, D. An electrophysiological investigation into the pain-relieving effects of heterotopic nociceptive stimuli: Probable involvement of a supraspinal loop. Brain 110, 1497–1508 (1987).

3. Villanueva, L., Cadden, S. W. & Le Bars, D. Evidence that diffuse noxious inhibitory controls (DNIC) are medicated by a final post-synaptic inhibitory mechanism. Brain Res. 298, 67–74 (1984).

4. Bannister, K., Patel, R., Goncalves, L., Townson, L. & Dickenson, A. H. Diffuse noxious inhibitory controls and nerve injury: restoring an imbalance between descending monoamine inhibitions and facilitations. Pain 156, 1803–11 (2015).

5. Dickenson, A. H. & Le Bars, D. Diffuse noxious inhibitory controls (DNIC) involve trigeminothalamic and spinothalamic neurones in the rat. Exp. brain Res. 49, 174–80 (1983).

6. De Broucker, T., Cesaro, P., Willer, J. C. & Le Bars, D. Diffuse noxious inhibitory controls in man. Involvement of the spinoreticular tract. Brain 113 (Pt 4, 1223–34 (1990).

7. Bouhassira, D., Bing, Z. & Le Bars, D. Effects of lesions of locus coeruleus/subcoeruleus on diffuse noxious inhibitory controls in the rat. Brain Research 571, 140–144 (1992).

8. Bouhassira, D., Bing, Z. & Le Bars, D. Studies of the brain structures involved in diffuse noxious inhibitory controls: the mesencephalon. J. Neurophysiol. 64, 1712–23 (1990).

9. Bouhassira, D., Villanueva, L., Bing, Z. & le Bars, D. Involvement of the subnucleus reticularis dorsalis in diffuse noxious inhibitory controls in the rat. Brain Res. 595, 353–7 (1992).

10. Barik, A., Thompson, J. H., Seltzer, M., Ghitani, N. & Chesler, A. T. A Brainstem-Spinal Circuit Controlling Nocifensive Behavior. Neuron 100, 1491-1503.e3 (2018).

11. Bannister, K., Lockwood, S., Goncalves, L., Patel, R. & Dickenson, A. H. H. An investigation into the inhibitory function of serotonin in diffuse noxious inhibitory controls in the neuropathic rat. Eur. J. Pain (United Kingdom) 21, 750–760 (2017).

12. Kucharczyk, M. W., Di Domenico, F. & Bannister, K. Distinct brainstem to spinal cord noradrenergic pathways inversely regulate spinal neuronal activity. Brain 1–18 (2022). doi:10.1093/brain/awac085

13. Lyons, W. E., Fritschy, J. M. & Grzanna, R. The noradrenergic neurotoxin DSP-4 eliminates the coeruleospinal projection but spares projections of the A5 and A7 groups to the ventral horn of the rat spinal cord. J. Neurosci. 9, 1481–9 (1989).

14. Fritschy, J.-M & Grzanna, R. Demonstration of two separate descending noradrenergic pathways to the rat spinal cord: Evidence for an intragriseal trajectory of locus coeruleus axons in the superficial layers of the dorsal horn. J. Comp. Neurol. 291, 553–582 (1990).

15. Tavares, I. The pontine A5 noradrenergic cells which project to the spinal cord dorsal horn are reciprocally connected with the caudal ventrolateral medulla in the rat. European Journal of Neuroscience 9, 2452–2461 (1997).

16. Bruinstroop, E. et al. Spinal projections of the A5, A6 (locus coeruleus), and A7 noradrenergic cell groups in rats. J. Comp. Neurol. 520, 1985–2001 (2012).

17. Villanueva, L., Cadden, S. W. & Le Bars, D. Diffuse noxious inhibitory controls (DNIC): evidence for post-synaptic inhibition of trigeminal nucleus caudalis convergent neurones. Brain Res. 321, 165–8 (1984).

18. Phelps, C. E., Navratilova, E., Dickenson, A. H., Porreca, F. & Bannister, K. Kappa opioid signaling in the right central amygdala causes hind paw specific loss of diffuse noxious inhibitory controls in experimental neuropathic pain. Pain 160, 1614–1621 (2019).

19. Morgan, M. M., Gogas, K. R. & Basbaum, A. I. Diffuse noxious inhibitory controls reduce the expression of noxious stimulus-evoked Fos-like immunoreactivity in the superficial and deep laminae of the rat spinal cord. Pain 56, 347–52 (1994).

20. Wen, Y.-R. et al. DNIC-mediated analgesia produced by a supramaximal electrical or a high-dose formalin conditioning stimulus: roles of opioid and alpha2-adrenergic receptors. J. Biomed. Sci. 17, 19 (2010).

21. Kucharczyk, M. W., Valiente, D. & Bannister, K. Developments in Understanding Diffuse Noxious Inhibitory Controls: Pharmacological Evidence from Pre-Clinical Research. J. Pain Res. 14, 1083–1095 (2021).

22. Pertovaara, A., Haapalinna, A., Sirviö, J. & Virtanen, R. Pharmacological properties, central nervous system effects, and potential therapeutic applications of atipamezole, a selective α2-adrenoceptor antagonist. CNS Drug Rev. 11, 273–288 (2005).

23. Westlund, K. N., Bowker, R. M., Ziegler, M. G. & Coulter, J. D. Noradrenergic projections to the spinal cord of the rat. Brain Res. 263, 15–31 (1983).

24. Li, Y. et al. Retrograde optogenetic characterization of the pontospinal module of the locus coeruleus with a canine adenoviral vector. Brain Res. 1641, 274–290 (2016).

25. Villanueva, L., Chitour, D. & Le Bars, D. Involvement of the dorsolateral funiculus in the descending spinal projections responsible for diffuse noxious inhibitory controls in the rat. J. Neurophysiol. 56, 1185–95 (1986).

26. Treweek, J. B. et al. Whole-body tissue stabilization and selective extractions via tissue-hydrogel hybrids for high-resolution intact circuit mapping and phenotyping. Nat. Protoc. 10, 1860–1896 (2015).

27. Clark, F. M. & Proudfit, H. K. The projections of noradrenergic neurons in the A5 catecholamine cell group to the spinal cord in the rat: anatomical evidence that A5 neurons modulate nociception. Brain Res. 616, 200–210 (1993).

28. Clark, F. M. & Proudfit, H. K. The projection of locus coeruleus neurons to the spinal cord in the rat determined by anterograde tracing combined with immunocytochemistry. Brain Res. 538, 231–245 (1991).

29. Clark, F. M. & Proudfit, H. K. The projection of noradrenergic neurons in the A7 catecholamine cell group to the spinal cord in the rat demonstrated by anterograde tracing combined with immunocytochemistry. Brain Res. 547, 279–88 (1991).

30. Mahn, M. et al. High-efficiency optogenetic silencing with soma-targeted anion-conducting channelrhodopsins. Nat. Commun. 9, (2018).

31. Hayat, H. et al. Locus coeruleus norepinephrine activity mediates sensory-evoked awakenings from sleep. Sci. Adv. 6, (2020).

32. Mahn, M., Prigge, M., Ron, S., Levy, R. & Yizhar, O. Biophysical constraints of optogenetic inhibition at presynaptic terminals. Nat. Neurosci. 19, 554–556 (2016).

33. Tervo, D. G. R. et al. A Designer AAV Variant Permits Efficient Retrograde Access to Projection Neurons. Neuron 92, 372–382 (2016).

34. Chuong, A. S. et al. Noninvasive optical inhibition with a red-shifted microbial rhodopsin. Nat. Neurosci. 17, 1123–9 (2014).

35. Ganley, R. P., Werder, K., Wildner, H. & Zeilhofer, H. U. Spinally projecting noradrenergic neurons of the locus coeruleus display resistance to AAV2retro-mediated transduction. Mol. Pain 17, (2021).

36. Hickey, L. et al. Optoactivation of Locus Ceruleus Neurons Evokes Bidirectional Changes in Thermal Nociception in Rats. J. Neurosci. 34, 4148–4160 (2014).

37. Li, Y. et al. Retrograde optogenetic characterization of the pontospinal module of the locus coeruleus with a canine adenoviral vector. Brain Res. 1641, 274–290 (2016).

38. Bannister, K., Lockwood, S., Goncalves, L., Patel, R. & Dickenson, A. H. An investigation into the inhibitory function of serotonin in diffuse noxious inhibitory controls in the neuropathic rat. Eur. J. Pain 21, 750–760 (2017).

39. Pertovaara, A. The noradrenergic pain regulation system: a potential target for pain therapy. Eur. J. Pharmacol. 716, 2–7 (2013).

40. Abraira, V. E. et al. The Cellular and Synaptic Architecture of the Mechanosensory Dorsal Horn. Cell 168, 295-310.e19 (2017).

41. Usoskin, D. et al. Unbiased classification of sensory neuron types by large-scale single-cell RNA sequencing. Nat. Neurosci. 18, 145–153 (2014).

42. Todd, A. J. Neuronal circuitry for pain processing in the dorsal horn. Nat. Rev. Neurosci. 11, 823–36 (2010).

43. Rajaofetra, N., Poulat, P., Marlier, L., Geffard, M. & Privat, A. Pre- and postnatal development of noradrenergic projections to the rat spinal cord: an immunocytochemical study. Dev. Brain Res. 67, 237–246 (1992).

44. Zoli, M. & Agnati, L. F. Wiring and volume transmission in the central nervous system: The concept of closed and open synapses. Prog. Neurobiol. 49, 363–380 (1996).

45. Pertovaara, A. Noradrenergic pain modulation. Prog. Neurobiol. 80, 53–83 (2006).

46. Okada-Ogawa, A., Porreca, F. & Meng, I. D. Sustained morphine-induced sensitization and loss of diffuse noxious inhibitory controls in dura-sensitive medullary dorsal horn neurons. J. Neurosci. 29, 15828–35 (2009).

47. Lapirot, O. et al. NK1 receptor-expressing spinoparabrachial neurons trigger diffuse noxious inhibitory controls through lateral parabrachial activation in the male rat. Pain 142, 245–254 (2009).

48. Bester, H., Chapman, V., Besson, J. M. & Bernard, J. F. Physiological properties of the lamina I spinoparabrachial neurons in the rat. J. Neurophysiol. 83, 2239–59 (2000).

49. Suzuki, R., Morcuende, S., Webber, M., Hunt, S. P. & Dickenson, A. H. Superficial NK1-expressing neurons control spinal excitability through activation of descending pathways. Nat. Neurosci. 5, 1319–1326 (2002).

50. Fields, H. L., Heinricher, M. M. & Mason, P. Neurotransmitters in nociceptive modulatory circuits. Annu. Rev. Neurosci. 14, 219–45 (1991).

51. Millan, M. J. Descending control of pain. Progress in Neurobiology 66, 355–474 (2002).

52. Wei, H. & Pertovaara, A. Spinal and pontine alpha2-adrenoceptors have opposite effects on pain-related behavior in the neuropathic rat. Eur. J. Pharmacol. 551, 41–9 (2006).

53. Pereira-Silva, R. et al. Attenuation of the Diffuse Noxious Inhibitory Controls in Chronic Joint Inflammatory Pain Is Accompanied by Anxiodepressive-Like Behaviors and Impairment of the Descending Noradrenergic Modulation. Int. J. Mol. Sci. 21, 1–28 (2020).

54. Kucharczyk, M. W., Derrien, D., Dickenson, A. H. & Bannister, K. The Stage-Specific Plasticity of Descending Modulatory Controls in a Rodent Model of Cancer-Induced Bone Pain. Cancers (Basel). 12, 1–17 (2020).

55. Villanueva, L., Bernard, J. F. & Le Bars, D. Distribution of spinal cord projections from the medullary subnucleus reticularis dorsalis and the adjacent cuneate nucleus: A phaseolus vulgaris-leucoagglutinin study in the rat. J. Comp. Neurol. 352, 11–32 (1995).

56. Bannister, K., Kucharczyk, M. W., Graven-Nielsen, T. & Porreca, F. Introducing descending control of nociception: a measure of diffuse noxious inhibitory controls in conscious animals. Pain Publish Ah, (2021).

57. Da Silva, J. T., Zhang, Y., Asgar, J., Ro, J. Y. & Seminowicz, D. A. Diffuse noxious inhibitory controls and brain networks are modulated in a testosterone-dependent manner in Sprague Dawley rats. Behav. Brain Res. 349, 91–97 (2018).

58. Da Silva, J. T., Tricou, C., Zhang, Y., Seminowicz, D. A. & Ro, J. Y. Brain networks and endogenous pain inhibition are modulated by age and sex in healthy rats. Pain 00, 1 (2020).

59. Yarnitsky, D. Conditioned pain modulation (the diffuse noxious inhibitory control-like effect): Its relevance for acute and chronic pain states. Curr. Opin. Anaesthesiol. 23, 611–615 (2010).

60. Yarnitsky, D. et al. Recommendations on terminology and practice of psychophysical DNIC testing. Eur. J. Pain 14, 339 (2010).

61. Niesters, M. et al. Tapentadol potentiates descending pain inhibition in chronic pain patients with diabetic polyneuropathy. Br. J. Anaesth. 113, 148–56 (2014).

62. Yarnitsky, D. Conditioned pain modulation (the diffuse noxious inhibitory control-like effect): Its relevance for acute and chronic pain states. Curr. Opin. Anaesthesiol. 23, 611–615 (2010).

63. Yarnitsky, D., Granot, M., Nahman-Averbuch, H., Khamaisi, M. & Granovsky, Y. Conditioned pain modulation predicts duloxetine efficacy in painful diabetic neuropathy. Pain 153, 1193–1198 (2012).

64. Lockwood, S. M., Lopes, D. M., McMahon, S. B. & Dickenson, A. H. Characterisation of peripheral and central components of the rat monoiodoacetate model of Osteoarthritis. Osteoarthr. Cartil. 27, 712–722 (2019).

65. Youssef, A. M., Macefield, V. G. & Henderson, L. A. Cortical influences on brainstem circuitry responsible for conditioned pain modulation in humans. Hum. Brain Mapp. 37, 2630–2644 (2016).

66. Zimmermann, M. Ethical guidelines for investigations of experimental pain in conscious animals. Pain 16, 109–10 (1983).

67. Kilkenny, C., Browne, W. J., Cuthill, I. C., Emerson, M. & Altman, D. G. Improving bioscience research reporting: the ARRIVE guidelines for reporting animal research. PLoS Biol. 8, e1000412 (2010).

68. Krashes, M. J. et al. Rapid, reversible activation of AgRP neurons drdives feeding behavior in mice. J. Clin. Invest. 121, 1424–28 (2011).

69. Urch, C. E. & Dickenson, a. H. In vivo single unit extracellular recordings from spinal cord neurones of rats. Brain Res. Protoc. 12, 26–34 (2003).

70. LeChasseur, Y. et al. A microprobe for parallel optical and electrical recordings from single neurons in vivo SUPPLEMENT. Nat Methods 8, 319–325 (2011).

