## Supplementary figures and images for "The origin of diffuse noxious inhibitory controls"

### Figure S1

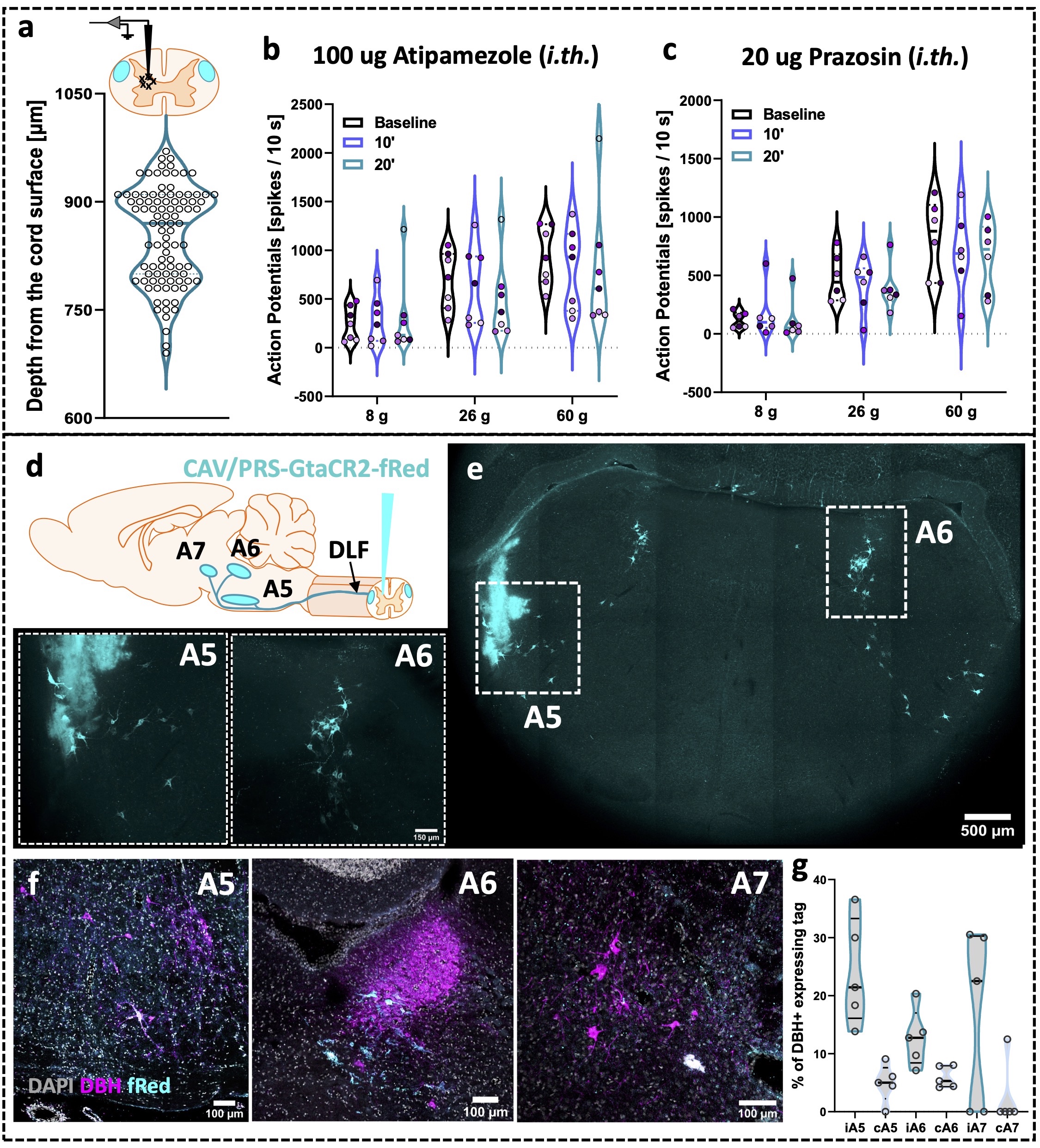

### Figure S2

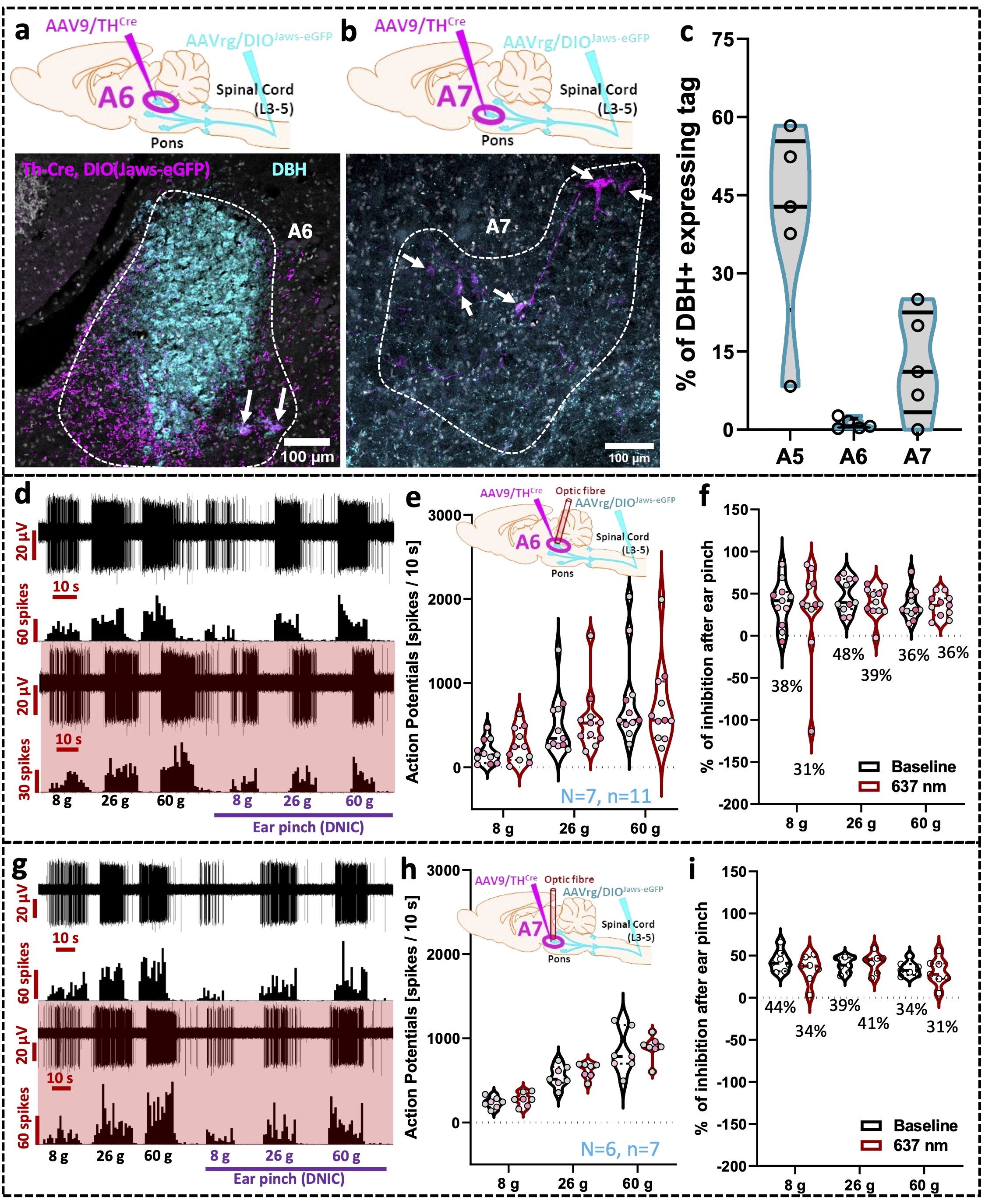

### Figure S3

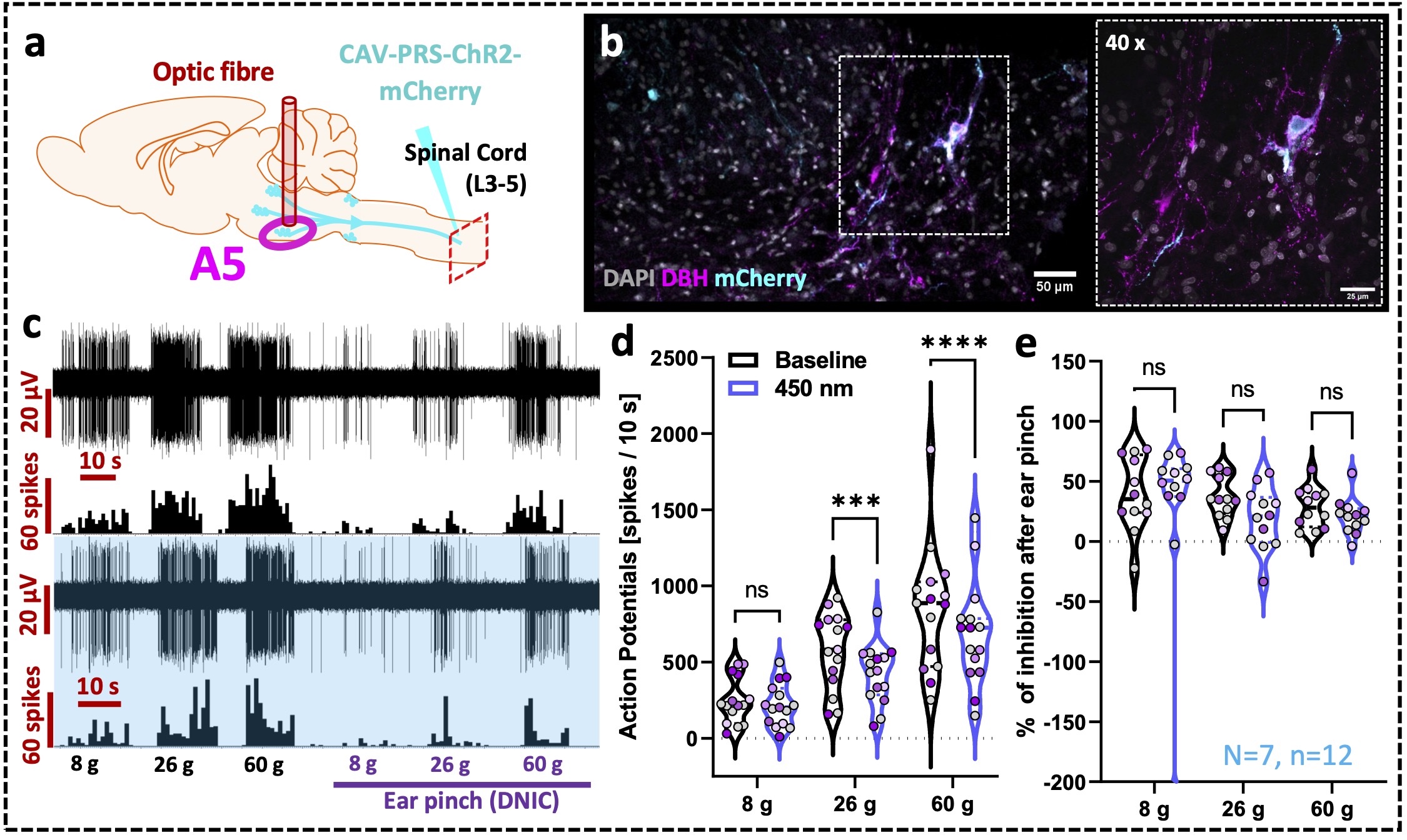
